# Domain-Specific Functional Plasticity of Visual Processing Constrained by General Cognitive Ability in Deaf Individuals

**DOI:** 10.64898/2026.03.25.714101

**Authors:** Chenjie Dong, Zhengye Wang, Xiaoyi Zuo, Suiping Wang

## Abstract

Interpersonal communication relies on integrating facial and vocal signals to extract multidimensional communicative information. How the absence of audition reshapes the communicative system remains unclear. We compared the performance of deaf (*N=136*) and hearing (*N=135*) adults across multiple domains, facial identity, emotional expression, speech, and global motion, through a series of unisensory and audiovisual psychophysical tasks. The results showed that, in hearing individuals, reliance on facial versus vocal signals differed across domains. In deaf individuals, auditory deprivation did not produce uniform enhancement or impairment of visual processing. Instead, they exhibited reduced sensitivity to dynamic emotional expressions and global motion, preserved sensitivity to facial identity (both static and dynamic) and static expressions, and enhanced categorization of facial speech. Notably, sensitivity to dynamic facial expressions and global motion was correlated, and both were explained by variations in fluid intelligence. Our results reveal a domain specific framework of functional plasticity in deaf individuals, suggesting that the consequences of hearing loss are shaped both by the functional roles of audition within each domain and by broader cognitive adaptations. This framework extends the classical deficit□or□compensation perspective, deepens understanding of cognitive plasticity, and informs the development of targeted, ecologically valid accessibility and sensory□substitution strategies.

## 1. Introduction

Efficient interpersonal communication relies on the seamless decoding of multidimensional communicative information, such as personal identity, emotional state, and speech, from concurrent facial and vocal signals^1–3^. In neurotypical individuals, these signals are integrated to optimize the perceptual inference during communication^4^. In contrast, individuals with sensory deprivation rely exclusively on intact modalities, leading to distinct patterns of perceptual inference^5^. This raises a central question in cognitive neuroscience: how does the absence of one sensory modality reshape processing of communicative information through remaining modalities? People with hearing loss provide a powerful model for addressing this question. In particular, how the visual system adapts to perceiving communicative information, such as emotional state, that is typically distributed across auditory and visual modalities^6–8^.

Two classical perspectives have been debated for several decades^9,10^. The functional deficit perspective proposes that auditory inputs are necessary for calibrating specific aspects of visual function, especially those requiring fine-grained temporal processing^11^. Consequently, auditory deprivation impair temporally demanding visual functions, while leaving fundamental visual functions intact^12,13^. Consistent with this view, deaf individuals show preserved basic visual functions, such as brightness perception^14^, color discrimination^15^, and facial configuration processing^16^, but reduction in visual temporal perception^12^, dynamic facial expression recognition^17,18^, and working memory^19^. Moreover, auditory speech deprivation leads to delayed language development^20^ and broader communication difficulties^21,22^.

In contrast, the functional compensation perspective suggests that auditory deprivation induces cross-modal reorganization in auditory cortices^5,23^, which, together with extensive visual training experience^12,24^, enhances a broad range of visual functions. Supporting this, neuroimaging studies have demonstrated recruitment of auditory cortical regions for visual processing in deaf individuals^25,26^. Notably, the vocal identity area in superior temporal cortex (STC) was engaged in facial identity processing in deaf individuals^27^. Behavioral studies revealed superior performance in motion perception^28,29^, facial identity recognition^30,31^, expression perception^32,33^, and lip-reading^34,35^ in deaf individuals. Furthermore, sign language experience has been associated with enhanced performance across several of these domains ^30,36^.

Although these two hypotheses appear to make opposing predictions, they are not theoretically incompatible. Indeed, systematic reviews^10,12,37^ and meta-analyses^38,39^ reveal a heterogeneous pattern of visual performance in deaf individuals. Low-level visual functions tend to remain stable, whereas temporally demanding or linguistically grounded processes often show deficits, and visually trained functions are frequently enhanced^10,12,37^. Neuroimaging evidence also demonstrated domain-general visual recruitment of the right STC associated with auditory deprivation, alongside language-related reorganization of left STC shaped by linguistic experience^40,41^. These findings converge on a domain□specific functional plasticity framework, in which sensory deprivation induces selective, rather than uniform modifications of visual funcation^9,37,42^.

However, whether this domain□specific plasticity is confined to perceptual□level visual functions or reflects a more general principle that extends to higher□order visual processes engaged in daily communication remains unclear. Specifically, how does the visual system of deaf individuals adapt to support the recognition of identity, emotional expressions, and speech, functions that rely on auditory and visual inputs to different degrees? Prior studies have typically examined isolated domains^18,30,35^, employed heterogeneous task demands, or relied on small sample sizes, limiting direct cross□domain comparisons and generalizability^38^.

To address these gaps, we examined how auditory deprivation shapes visual processing of communicative information by systematically comparing deaf and hearing adults across three domains, including facial identity, emotional expression, and speech. We expected that auditory deprivation would exert selective effects across domains. From the multisensory calibration hypothesis^60,61^, deaf individuals should show preserved facial identity perception, which relies comparably on facial and vocal signals, but reductions in emotional expressions processing, which require calibration of temporal variations from auditory signals, as well as reduced speech processing, which depends on calibration from auditory speech inputs. From the functional compensation hypothesis, deaf individuals should exhibit enhanced performance across domains, with the strongest gains in speech perception, followed by emotional expression, and then facial identity, with these enhancements linked to sign language and lip□reading experience.

To test these predictions, we designed a series of two□alternative forced□choice (2AFC) tasks examining categorization sensitivity in auditory, visual, and audiovisual modality. Deaf participants completed seven visual□only sessions, whereas hearing participants completed thirty sessions spanning three modalities. For facial identity and facial emotional expression, both static and dynamic facial stimuli were used to determine whether perceptual sensitivity depends on the temporal structure of facial information. For facial speech perception, participants viewed normal and temporally reversed articulations to dissociate sensitivity to articulatory structure from general motion cues. In addition, a visual global□motion task was included to assess whether the observed effects in the communication domain reflect broader alterations in the fundamental visual processes responsible for spatiotemporal signal integration.

## 2. Results

Across all two□alternative forced choice (2AFC) tasks, stimulus intensity was parametrically varied from 0% to 100% in 10% increments (Figure 1). Response proportions for the target category (e.g., happy) were fitted with cumulative Gaussian function (Figure S1-S2), yielding two parameters: slope, indexing perceptual sensitivity to variations in stimulus intensity, and threshold, reflecting categorical decision boundaries. This psychometric approach enabled comparisons of behavioral performance beyond accuracy on prototypical stimuli.

**Figure 1.**
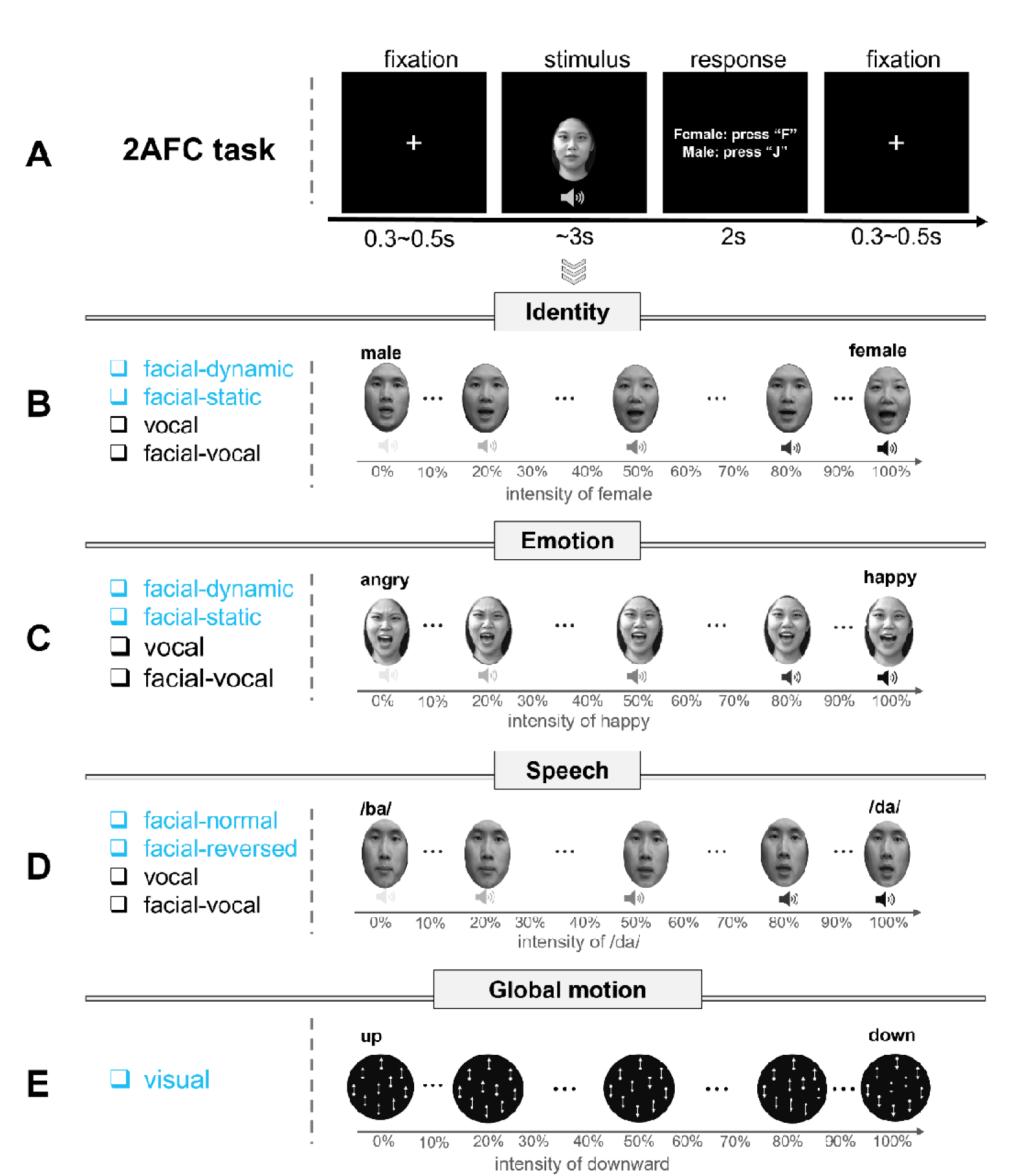
Experimental design and procedure. (A) Procedure of the two alternative forced choice (2AFC) tasks. (B) The identity task included four sessions - facial dynamic, facial static, vocal, and facial-vocal sessions - with identity intensity varying from 0% male features to 100% male features in 10% increments. (C) The emotion task included four sessions - facial dynamic, facial static, vocal, and facial-vocal sessions - with emotion intensity varying from 0% happy features to 100% happy features in 10% increments. (D) The speech task included four sessions – normal facial, reversed facial, vocal, and facial-vocal sessions - with speech intensity varying from 0% /ba/ syllable to 100% /ba/ syllable in 10% increments. (E) The global motion direction task with motion intensit varying from 0% downward to 100% downward in 10% increments. The deaf group only completed visual versio of tasks (in blue color). All videos used in the experiments were recordings of lab members who consented to the use of their images in the manuscript.

First, to characterize the contribution of auditory and visual modalities across communicative domains, we examined the performance (accuracy, perceptual sensitivity, and categorical boundaries) to facial signals, vocal signals, as well as their combination in identity, emotion, and speech tasks in hearing participants. Then we compared the performance of deaf and hearing participants to assess how auditory deprivation reshapes visual performance across these domains that rely differently on facial and vocal signals. Specifically, we tested whether categorization accuracy, perceptual sensitivity, and categorical boundaries are uniformly affected or selectively altered using linear mixed effect (LME) model. After that, we predicted the perceptual sensitivity and categorical boundaries by demographic factors (e.g., age) and domain□general cognitive ability (Raven’s Standard Progressive Matrices scores) using generalized linear model (GLM). Finally, we conducted four control analyses to assess the robustness of the results.

### 2.1 Participant Demographics

Detailed demographic information is provided in Table□S6. Deaf participants were significantly older than the hearing participants (*t*_(236)_ =-10.39; *p* = 4.57×10^-21^) and exhibited lower Raven’s scores (*t*_(267)_ = 7.48; *p* = 1.08×10^-12^; Table 1).

**Table 1.** Demographics of the deaf and hearing participants.

|  | Hearing controls<br>(*N* = 135) | Deaf participants<br>(*N* = 136) | T value | *p*-value |
| --- | --- | --- | --- | --- |
| **Gender: males/females (n)** | 54/81 | 74/61 | / | / |
| **Age (years)** | 21.5 ± 2.2 | 24.1 ± 3.5 | -7.48 | < 0.001 |
| **RSPM Score** | 109.5 ± 12.5 | 92.3 ± 13.0 | 10.39 | < 0.001 |
| **Handedness: right/left (n)** | 132/2 | 116/19 | / | / |
| **Age of HL onset (years)** | / | 1.6 ± 2.1 | / | / |
| **Severe-to-profound HL (%)** | / | 97.1 | / | / |
| **Sign language users (%)** | / | 94.1 | / | / |
| **Age of SL onset (years)** | / | 9.1 ± 3.4 | / | / |
| **Proficiency of SL (1-10)** | / | 6.3 ± 1.7 | / | / |
| **Proficiency of lip-reading (1-5)** |  | 2.6 ± 0.9 |  |  |

RSPM = Raven's Standard Progressive Matrices; HL = hearing loss; SL = sign language; Self-reported SL proficiency: 1 = no ability, 10 = native/near-native fluency; Self-reported lip-reading proficiency: 1 = cannot understand any conversations, 5 = can understand most conversations; "/" not applicable to this group.

### 2.2 Facial-vocal facilitation across all communicative domains

In hearing group, facial and vocal signals contributed differentially across communicative domains (supplementary materials, Table S1-S2). For identity (Figure S3A), facial and vocal signals were comparable in accuracy, with a modest advantage for facial signal in perceptual sensitivity. For emotion (Figure S3B), facial signals were more reliable than vocal signals, whereas, for speech (Figure S3C), vocal signals were more reliable. Importantly, audiovisual integration enhanced perceptual sensitivity relative to unisensory conditions in all domains (Figure S3). These findings demonstrate that the relative reliance on facial and vocal signals differs across domains, providing a benchmark for assessing domain□specific changes of visual processing in the absence of auditory input.

### 2.3 Preserved facial identity perception in the absence of auditory input

We first examined whether auditory deprivation affects the perception of static and dynamic identity in vision, the marginally dominant modality for this process. For categorization accuracy on prototypical face (i.e., 90% male and 90% female, Table S3), the LME analysis revealed no significant main effects of group (deaf vs. hearing), stimulus type (static vs. dynamic), or their interaction.

For perceptual sensitivity (i.e., slope; Figure 2A, Table S4), LME analysis indicated a significant main effect of stimulus type (*F* _(1, 167)_ = 15.47, *p* = 1×10^-4^), with higher sensitivity for static than dynamic stimuli, no significant main effect of group, and no group × stimulus type interaction. Similarly, categorization boundaries (i.e., threshold; Figure 2A, Table S4) differed between static and dynamic face (*F* _(1, 167)_ = 7.831, *p* = 0.006) but did not differ between groups. These results indicate that facial identity processing remains intact in deaf individuals, regardless of facial movements.

**Figure 2.**
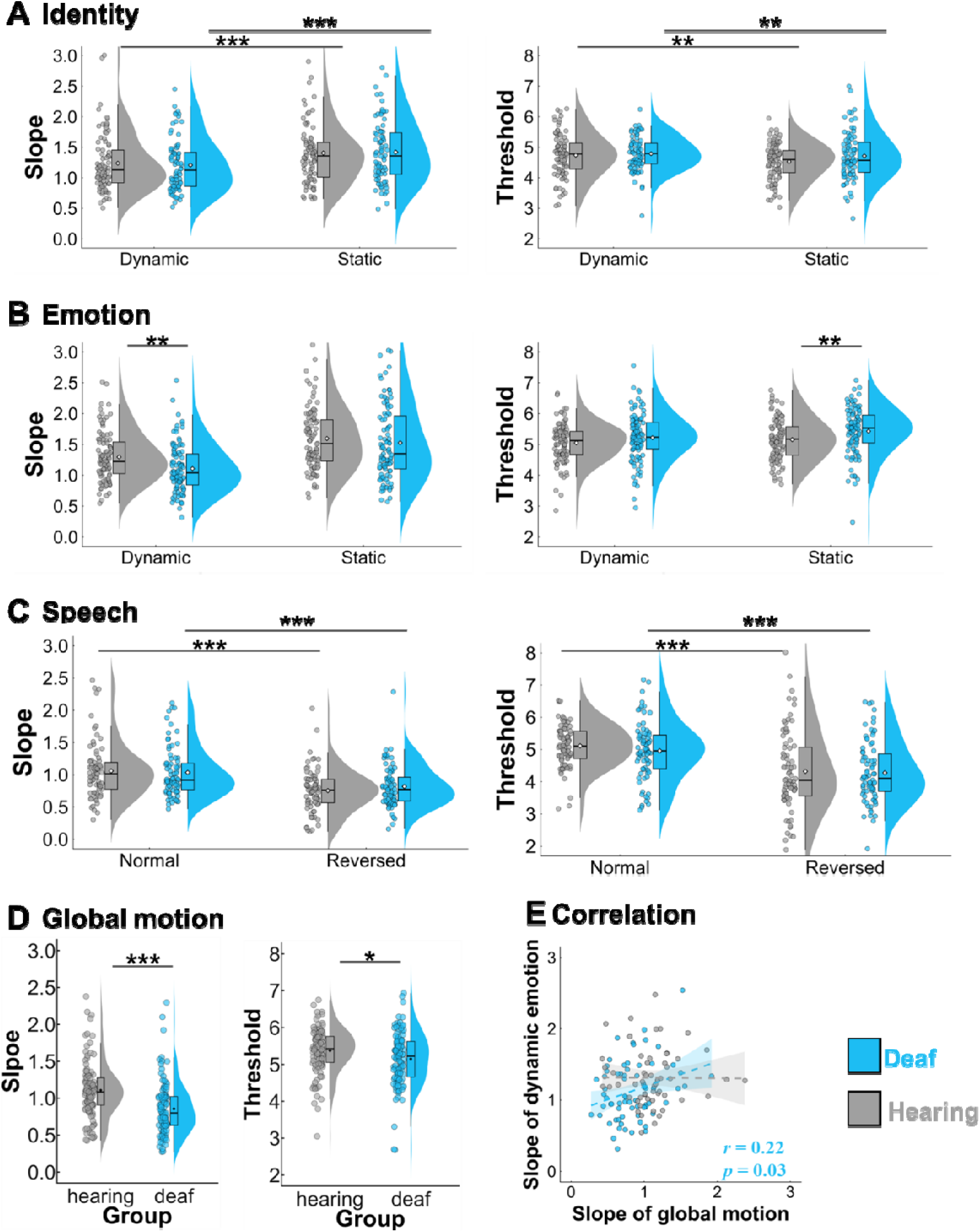
Psychometric parameters across visual tasks in the hearing and deaf groups. (A) Perceptual sensitivity (indexed by slope) and categorization boundaries (indexed by threshold) for facial identity were comparable between two groups, irrespective of stimulus type. (B) Perceptual sensitivity for dynamic facial expressions was reduced in the deaf group compared with the hearing group. (C) Perceptual sensitivity an categorization boundaries for facial speech were comparable between the two groups, irrespective of stimulus type. (D) Perceptual sensitivity for global motion was reduced in the deaf group compared with the hearing group. (E) Correlation between perceptual sensitivity in the global motion categorization and in the dynamic facial expression categorization.

### 2.4 Reduced sensitivity selectively for dynamic expression in the absence of auditory input

We next examined whether auditory deprivation affects the perception of emotional facial expressions, a domain that relies predominantly on visual modality and contains rich spatiotemporally varied features. For accuracy on prototypical stimuli (i.e., 90% happy and 90% angry; Table S3), the LME model revealed significant main effect of group (*F* _(1, 243)_ = 9.82, *p* = 0.002) and stimulus type (*F* _(1, 243)_ = 9.5, *p* = 0.002), and their interaction (*F* _(1, 243)_ = 5.04, *p* = 0.03). Post-hoc comparisons showed that deaf participants exhibited slightly lower accuracy than hearing participants for the dynamic expression (deaf = 98.5%±2.5%, hearing = 99.3%±1.2%; *t* _(472)_ =-3.8, *p _Bonferroni_* = 1.61 × 10^-4^, CI _95%=_ [-1%,-0.3%]), whereas no group difference was observed for static expressions.

For perceptual sensitivity (Figure 2B, Table S4), the LME model revealed a significant main effect of group (*F* _(1, 199)_ =4.79, *p* = 0.03), with lower sensitivity for deaf than hearing participants, and a significant main effect of task (*F* _(1, 199)_ =84.14, *p* = 5.92 × 10^-17^), with lower sensitivity for dynamic compared to static stimuli. Although the group × stimulus type interaction did not reach significance, planned exploratory post-hoc analysis revealed that deaf participants showed reduced sensitivity specifically for dynamic expressions (deaf = 1.1±0.4, hearing =1.3±0.4; *t* _(347)_ =-2.66, *p_Bonferroni_*= 0.008, CI _95%=_ [-0.33, - 0.05]), but not for static expressions (deaf = 1.53±0.62, hearing = 1.6±0.54; *t* _(347)_ =-0.98, *p_Bonferroni_* = 0.33).

For the categorization boundaries (Figure 2B, Table S4), the model revealed main effects of group (*F* _(1, 199)_ = 6.0, *p* = 0.02) and stimulus type (*F* _(1, 199)_ = 11.42, *p_Bonferron_* = 8×10^-4^), but no significant interaction. Exploratory post-hoc comparisons indicated higher thresholds in deaf than hearing participants for static expressions (deaf = 5.43±0.76; hearing = 5.16±0.67; *t* _(298)_ = 2.69, *p_Bonferroni_*= 0.007, CI _95%=_ [0.07, 0.47]), suggesting a bias toward categorizing ambiguous expressions as “happy”. No significant group difference was observed for dynamic expressions (deaf = 5.22±0.78, hearing = 5.05±0.64).

These results suggest that auditory deprivation selectively reduces sensitivity to dynamic facial expressions, while leaving static expression perception preserved.

### 2.5 Preserved facial speech processing in the absence of auditory input

We then assessed whether reduced sensitivity to dynamic signals generalize to normal and temporally reversed facial speech, a domain in which vision is the non dominant modality. For categorization accuracy on prototypical syllable (i.e., 90% ba and 90% da; Table S3), the LME model revealed significant main effects of group (*F* _(1, 199)_ = 6.16, *p* = 0.014) and stimulus type (*F* _(1, 199)_ = 107.40, *p_Bonferroni_* = 2.12× 10^-20^), and their interaction (*F* _(1, 199)_ = 8.60, *p* = 0.004). Post-hoc comparisons indicated that deaf participants performed significantly better than hearing participants when categorizing reversed facial speech (deaf = 90.7%±8.7%, hearing = 86.4%±11.5%; *t* _(393)_ =-3.81, *p_Bonferroni_* = 1.64 × 10^-4^, CI _95%_= [2.1%, 6.5%]). No significant group difference was observed for normal facial speech (deaf = 96.3%±4.5%, hearing = 96.4% ± 4.6%).

For perceptual sensitivity (Figure 2C, Table S4), the model revealed a significant main effect of stimulus type (*F* _(1, 156)_ = 45.25, *p* = 3.1 × 10^-14^), with higher sensitivity for normal than reversed speech, but no main effect of group and no group × stimulus type interaction. Similarly, for category boundaries (Figure 2C, Table S4), the model revealed a significant main effect of stimulus type (*F* _(1,156)_ = 62.04, *p* = 5.36 × 10^-13^), with higher thresholds for normal than reversed speech, but no group differences or significant interactions.

Together, auditory deprivation does not impair the categorization of speech from facial signals. Notably, deaf participants outperformed hearing participants in accuracy for reversed facial speech, a challenging condition that disrupts the normal temporal structure of articulation.

### 2.6 Reduced sensitivity extends beyond facial expression to global motion

To determine whether the reduced sensitivity observed for dynamic facial expressions reflects a broader change in visual spatiotemporal integration, we examined performance in global motion perception. For the accuracy on prototypical stimuli (90% downward and 90% upward; Table S3), deaf participants exhibited lower accuracy than hearing participants (deaf = 94.8%±6.2%, hearing = 96.8% ± 4.3%; *t* _(238)_ =-2.9, *p* = 0.004, CI _95%=_ [-3%,-0.6%]), although performance in both groups remained near ceiling.

For the two psychometric parameters (Figure 2D, Table S4), deaf participants showed significantly reduced sensitivity to global motion direction (deaf = 0.86 ± 0.36, hearing = 1.11 ± 0.38; *t* =-5.09, *p* = 7.85 × 10^-7^, CI_95%_ = [-0.35,-0.15]) and lower categorization thresholds (deaf = 5.15±0.73, hearing = 5.37 ± 0.61; *t* =-2.46, *p* = 0.01, CI_95%_ = [-0.4,-0.04]), compared with hearing participants.

These results parallel the reduced sensitivity for dynamic facial expressions, suggesting that auditory deprivation may selectively alter computations supporting the integration of local spatiotemporal signals into coherent global representations, spanning from low□level motion perception to higher level visual processing of dynamic communicative information.

### 2.7 General cognitive ability accounts for individual differences in perceptual sensitivity

We next assessed which demographic and cognitive factors were associated with individual variability in task performance. We fitted generalized linear models (GLM) predicting sensitivity from age and Raven’s score in both groups, and additionally from age of hearing loss, signlJlanguage proficiency, and lip reading proficiency within the deaf group.

Within the deaf group, Raven’s score significantly predicted perceptual sensitivity across multiple visual tasks, including static facial identity (Figure 3A, β = 0.008, *t* = 2.72, *p* = 0.03), static facial expression (Figure 3B, β = 0.01, *t* = 2.77, *p* = 0.007), dynamic facial expression (Figure 3C, β = 0.008, *t* = 2.84, *p* = 0.006), and global motion categorization (Figure 3D, β = 0.01, *t* = 3.56, *p* = 6.2 × 10^-4^). For instance, a ten point increase in Raven’s score was associated with an approximate 8%-12% increase in sensitivity across these tasks. Age of hearing loss additionally predicted sensitivity in the global motion task (β = 0.06, *t* = 2.95, *p* = 0.004), with later onset associated with higher sensitivity. In contrast, age, sign□language proficiency, and lip reading proficiency did not significantly predict sensitivity in any task.

**Figure 3.**
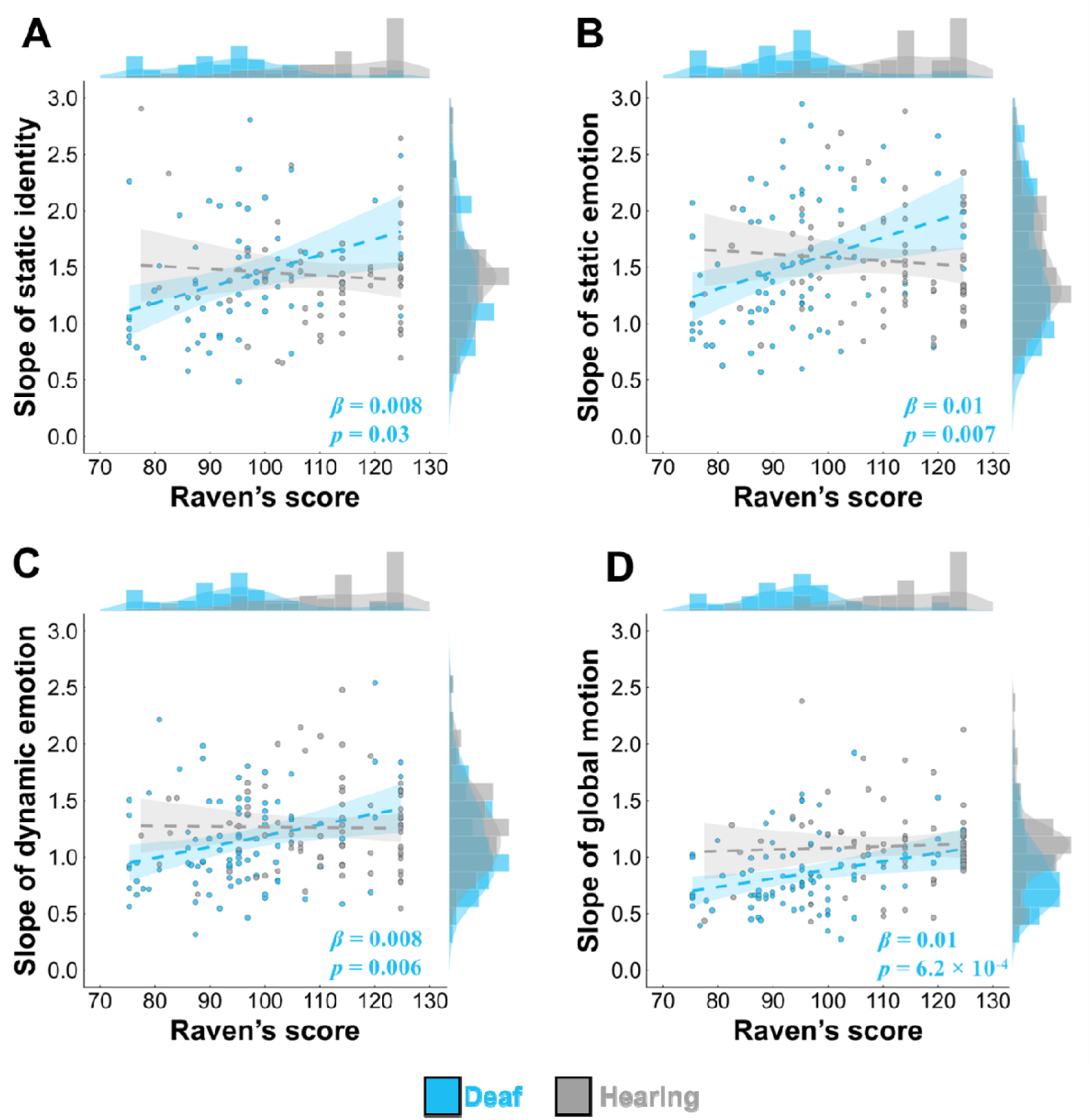
Raven’s score predicted individual differences of perceptual sensitivity only in deaf participants. (A) Predication in static facial identity categorization. (B) Predication in static facial emotion categorization. (C) Predication in dynamic facial emotion categorization. (D) Predication in global motion categorization.

Within the hearing group, age predicted sensitivity in dynamic facial identity categorization (β = 0.06*, t* = 0.02*, p* = 0.001), whereas Raven’s score did not predict sensitivity in any task.

Analyses of thresholds also indicated the association between Raven’s score and categorical biases (in supplementary results).

### 2.8 Control analyses

To assess the robustness of our findings, first, we examined cross□task correlations to evaluate whether reduced sensitivity reflected task□specific effects or shared perceptual processing. In deaf participants, perceptual sensitivity for dynamic facial expressions was significantly correlated with sensitivity for global motion (Figure 2E*, Spearman’s r* = 0.22, *p* = 0.03, *Fisher’s z* =0.23), whereas no such relationship was observed in hearing participants (Figure 2E, *Spearman’s r* = 0.005, *p* = 0.96). Paired-wise correlations among all tasks were provided in Figure S4.

Second, we conducted key analyses with age and Raven’s scores included as covariates and confirmed that reduced global motion sensitivity in deaf individuals persisted after accounting for individual differences in these variables (*t_(156)_* =-2.12, *p* = 0.04, CI_95%_ = [-0.31,-0.01]).

After that, we performed LME analysis using data from participants (number of deaf participants = 78; number of hearing participants = 71), who completed all four dynamic visual sessions (facial identity, emotional expression, normal speech, global motion).The model revealed a significant group × stimulus type interaction (*F* _(3, 441)_ = 3.7, *p* = 0.01). Compared with hearing participants, deaf participants exhibited reduced sensitivity for identity (*t_(568)_*=-2.87, *p* = 0.004, CI_95%_ = [-0.33,-0.06]), expression(*t_(568)_* =-3.72, *p* = 0.0002, CI_95%_ = [-0.39,-0.12]), and global motion(*t_(568)_* =-4.63, *p* = 4.47 × 10^-6^, CI_95%_ = [-0.45,-0.18]), but not for speech (*t_(568)_* =-0.41, *p* = 0.68).

Finally, we repeated key analyses in subgroups of deaf and hearing participants matched for age and Raven’s scores (Table S5). Deaf participants exhibited reduced sensitivity for dynamic facial expression categorization (Figure 4A, *t_(58)_* =-2.39, *p* = 0.02, CI_95%_ = [-1.14,-0.1]) and global motion categorization (Figure 4B*, t_(58)_* =-2.44, *p* = 0.02, CI_95%_ = [-1.15,-0.11]). In deaf participants, perceptual sensitivity for dynamic facial expressions was significantly correlated with sensitivity for global motion (Figure 4C*, Spearman’s r* = 0.35, *p* = 0.003, *Fisher’s z* =0.23) and Raven’s scores (Figure 4D*, Spearman’s r* = 0.42, *p* = 0.03, *Fisher’s z* = 0.45). Additionally, deaf participants still outperformed hearing participants in accuracy for reversed facial speech (deaf = 93%±7%, hearing = 87%±12%; *t_(58)_* = 2.45, *p* = 0.02, CI_95%_ = [11%, 15%]). These control analyses confirm that the observed reductions in perceptual sensitivity are robust, domainlJspecific, and not fully explained by demographic or general cognitive factors.

**Figure 4.**
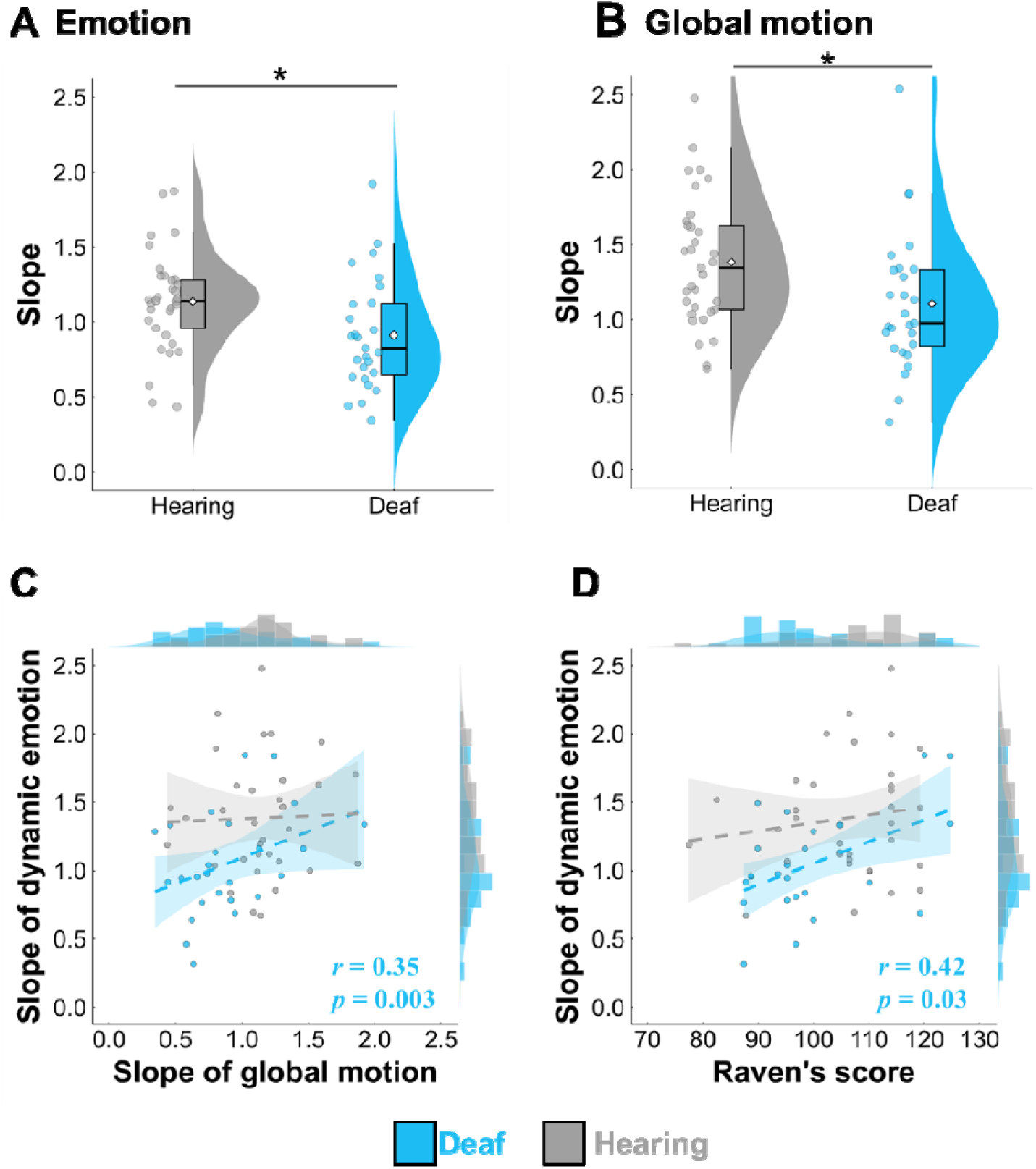
Psychometric parameters across visual tasks in hearing and deaf groups matched on demographic variables. (A) Perceptual sensitivity for dynamic facial expressions was reduced in the deaf grou compared with the hearing group. (B) Perceptual sensitivity for global motion was reduced in the deaf grou compared with the hearing group. (C) Correlation between perceptual sensitivity for the dynamic facial expressio and the global motion. (D) Correlations between Raven’s score and perceptual sensitivity for dynamic expressio in deaf group.

## 3. Discussion

Using the largest laboratory-based psychophysical dataset on individuals with hearing loss to date, we revealed domain□specific functional plasticity in the visual processing of deaf individuals. Specifically, the deaf group exhibited (i) reduced perceptual sensitivity to dynamic emotional expressions and global motion signals, (ii) preserved sensitivity to facial identity (both static and dynamic), static expressions, and normal facial articulations, and (iii) enhanced categorization of reversed facial articulations. Notably, sensitivity to dynamic expressions and global motion was correlated, and both were explained by variations in fluid intelligence, indexed by Raven’s matrices scores. Higher Raven’s scores were associated with increased perceptual sensitivity in these tasks. These findings suggest that auditory deprivation does not produce uniform enhancement or impairment of visual processing^9,43^. The functional consequences of hearing loss are shaped by both the role of audition in each domain and broader changes in the cognitive system.

Auditory deprivation does not uniformly alter facial information processing (identity, speech, emotion), but instead affects performance in specific domains. In the identity domain, the deaf group performed comparably to hearing controls^44–46^. Within the speech domain, the deaf group demonstrated equivalent performance to the hearing group^47,48^ and even exhibited an advantage for challenging facial articulations^34,35,49,50^. However, in the emotional expression domain, reduced sensitivity in deaf individuals emerged specifically for dynamic expressions, while perception of static expressions remained intact^51–53^. Similarly, studies using facial recognition paradigms reported lower recognition scores in deaf signers compared with hearing signers only for dynamic facial expressions, but not for static facial expressions^18^. These domain-specific dissociations^54,55^ align with the hierarchical processing model of face-voice perception, which proposes that identity, emotion, and speech involve distinct spatiotemporal features and are processed through functionally independent pathways^1^.

Notably, the reduced sensitivity observed in deaf group extended beyond dynamic emotional expressions and was also present in the global motion task. These decreases may reflect suboptimal visual categorization in deaf individuals. Supporting this, facial-vocal signals increased sensitivity in hearing controls relative to unisensory signals, suggesting that enhanced perceptual sensitivity reflects a more optimal strategy^56,57^. These decreases were not driven by changes in spatial (e.g., facial identity) or temporal processing (e.g., fluctuations in lip area) alone but may instead reflect altered spatiotemporal integration. For dynamic facial expressions, both the underlying spatial structure and the temporal trajectories of the signals must be consolidated^58,59^; similarly, the global motion task requires the construction of holistic representations by integrating the movements of spatially distributed dots. Moreover, the correlation between sensitivities to dynamic expressions and global motion, uniquely within the deaf group, suggests shared underlying processes in these tasks. These findings align with previous evidence suggesting that auditory deprivation may constrain the development of complex temporal representations in the visual domain^60,61^.

The observed reductions in visual spatiotemporal processing may indicate a functional trade off associated with reallocating attentional resources away from central and toward peripheral visual fields^38,62^. Deaf individuals exhibited broader spatial attention and heightened sensitivity to peripheral stimuli compared to hearing controls, reflecting compensatory adaptations to the absence of auditory input^63–65^. When signals were presented simultaneously at the center and periphery, deaf individuals were more susceptible to distraction from changes occurring in peripheral regions^63^. In dynamic facial expression tasks, hearing individuals benefited from anchoring their gaze on the center of the face (e.g., the nose), which facilitated efficient integration of motion cues across the entire face over time^59^. In contrast, deaf individuals showed broader and rapidly shifting attention across both relevant features and distractors, limiting the extraction of coherent facial information and reducing sensitivity in categorization. Overall, shifting attention toward the periphery enhances local detection and reorienting^63,66^, but leads to greater distractibility^64,67^ and reduced temporal allocation of attention required for global spatiotemporal integration^62,68^.

Crucially, our results point to broader consequences of auditory deprivation, extending from visual spatiotemporal processing to general cognitive abilities as indexed by lower Raven’s Progressive Matrices scores in the deaf group compared to the hearing group^69,70^. The large sample size enabled us to examine how demographic and cognitive factors relate to task performance. Notably, Raven scores were the only factor that positively predicted sensitivity to both dynamic emotional expressions and global motion in the deaf group. Analyses using age and Raven□matched subgroups confirmed that this association was not driven by group differences in age or nonverbal reasoning. Although the causal mechanisms underlying these relationships remain unclear, the findings suggest that deafness, and its cascading developmental consequences, affects multiple components of the cognitive system, including spatiotemporal processing (as shown here), executive functions (such as working memory^19^ and task switching^71^), and social interaction^72^. Whether these disruptions arise from auditory deprivation itself, delays in language acquisition^73,74^, or the inherent costs of extensive visual training^24^, as well as the neural mechanisms supporting these effects^43^, remains an open question.

In addition, from the perspective of multisensory integration, equivalent performance in facial identity and speech perception within the visual modality does not necessarily imply equivalent efficiency in real multisensory environments^55,75^. In our tasks, hearing participants optimized performance by integrating facial and vocal signals, gaining multisensory benefits^7^. In contrast, deaf participants relied solely on facial signals and therefore lacked these inherent audiovisual advantages, for example, they had to interpret ambiguous visual information without the clarifying support of tone of voice^76^. In typical multisensory contexts, this absence may place deaf individuals at a functional disadvantage, as achieving performance levels comparable to hearing peers likely requires greater compensatory effort and cognitive resources^77,78^. These findings underscore the importance of robust accessibility support and sensory-substitution strategies to promote more effective and equitable communication environments.

## 4. Conclusion

Auditory deprivation does not produce a uniform enhancement or impairment of visual abilities but instead gives rise to domain□specific functional plasticity. Whereas visual processing of facial identity and speech articulations were preserved or even enhanced, deaf individuals exhibited selective vulnerabilities in integrating spatiotemporal information, reflected in reduced sensitivity to dynamic emotional expressions and global motion. Sensitivities to these two domains were correlated and associated with individual differences in fluid intelligence. Our results suggest that the functional consequences of hearing loss are not only shaped by the role of audition in each domain but also by broader changes in the cognitive system. These findings extend the classical deficit□or□compensation perspective, deepens understanding of cognitive plasticity, and informs the development of targeted, ecologically valid accessibility and sensory□substitution strategies.

## 5. Methods

### 5.1 Participant

A total of one-hundred and thirty-six deaf adults (81 women; age = 24.1 ± 3.5 years) and one-hundred and thirty-five hearing controls (61 women; age = 21.5 ± 2.2 years) were recruited from local universities and community.

In the deaf group, participants were included only if they met criteria for severe-to-profound hearing loss. Exclusion criteria were i. pure-tone average threshold ≥ 71□dB HL (*n = 3*), ii. history of cochlear implantation (*n = 1*), and iii. voluntary withdrawal (*n = 1*).

All hearing controls reported normal audition and no prior experience with sign language.

Across both groups, all participants had normal or corrected-to-normal vision and reported no history of neurological or psychiatric disorders. Written informed consent was obtained from all participants prior to the experiment. They received monetary compensation for their participation.

### 5.2 Stimulus

#### 5.2.1 Identity stimulus

##### Prototypical dynamic identity

Two prototypical audiovisual identity stimuli were recorded from one male and one female speaker, each uttering the syllable /ba/. Video recordings were converted to grayscale and cropped with an elliptical mask to remove external features. During practice sessions, all participants achieved 100% accuracy in categorizing the prototypical facial and vocal identities.

##### Facial morphing

Nine intermediate facial identities were generated by interpolating features between the male and female prototypes in 10% increments. First, sixty-eight facial landmarks were extracted from the original videos, frame-by-frame, using the dlib library. Then, delaunay triangulation and affine transforms were applied to warp and blend the textures with OpenCV-2. This procedure yielded nine morphed faces (e.g., 10% male + 90% female features) that varied continuously along the identity continuum.

##### Vocal morphing

First, the original audio signals were decomposed into three parameters: fundamental frequency (F0), spectral envelope, aperiodicity. To generate the continuum, these parameters, along with the time and frequency axes, were mapped between the two voices to define five distinct morphing aspects. Weighted linear interpolation of these parameters was then used to generate nine vocal identities (e.g., 10% male + 90% female voice) that varied continuously along the identity continuum.

##### Static facial identity

Static stimuli were generated by extracting the first frame from each dynamic clip. These images were similarly converted to grayscale and masked to ensure consistency across static and dynamic stimuli. The static frame was repeated to create a video sequence with the exact same number of frames and total duration as the corresponding dynamic stimuli.

##### Audiovisual identity

Nine congruent audiovisual stimuli were constructed by pairing each facial morph with its corresponding vocal morph (e.g., 20% male face with 20% male voice) using Adobe Premiere.

The video clips were around 3 seconds long, recorded at a frame rate of 25 frames per second, with a resolution of 960 × 540 pixels. The audio stimuli were recorded at a sampling rate of 44100 kHz and presented at a sound level around 70 dB using noise-attenuating headphones.

#### 5.2.2 Expression stimulus

##### Prototypical dynamic expression

Two prototypical audiovisual emotion stimuli (happy and angry) were recorded from one female speaker, each uttering the syllable /a/. The video recordings were converted to grayscale and cropped with an elliptical mask to remove external features. Each clip depicted a natural unfolding of the expression from neutral onset to peak intensity. During practice sessions, all participants achieved 100% accuracy in categorizing the prototypical facial and vocal expressions.

##### Facial and vocal morphing

Nine intermediate facial and vocal expressions with10% increments were generated, using the prototypes (100% happy and 100% angry) as the morphing endpoints, followed the same procedures described for the identity stimuli.

##### Static facial stimuli

Static facial stimuli were generated by extracting the apex frame from each dynamic clip. The static apex frame was repeated to create a video sequence with the exact same number of frames and total duration as the corresponding dynamic expression stimuli.

The audiovisual synthesis and stimulus parameters were identical to those used for the identity stimuli.

#### 5.2.3 Speech stimulus

##### Prototypical normal speech

Two prototypical audiovisual speech stimuli (/ba/ and /da/) were recorded from one male speaker used in the identity task. The video recordings were converted to grayscale and cropped with an elliptical mask to remove external features. Each clip captured the natural articulatory progression from a neutral closed-mouth onset, peak of the vowel production, and closed-mouth offset. During practice sessions, all participants achieved 100% accuracy in categorizing the prototypical facial and vocal syllables.

##### Facial and vocal morphing

Nine intermediate facial and vocal speech with10% increments were generated, using the prototypical speech (100% /ba/ and 100% /da/) as the morphing endpoints, followed the same procedures described for the identity stimuli.

##### Reversed facial speech

The reversed facial speech stimuli were created by inverting the temporal sequence of the video frames from the normal speech clips (playing them from offset to onset), while maintaining the original grayscale and masking parameters.

The audiovisual synthesis and stimulus parameters were identical to those used for the identity stimuli.

#### 5.2.4 Global motion stimulus

##### Prototypical global motion stimuli

Two prototypical global motion stimuli were generated using random dot kinematogram. Each stimulus consisted of 100 white dots (dot size = 15 pixels; speed = 1.5 pixels/frame; dot lifetime = 80 frames) moving coherently within a circular aperture (diameter = 381 pixels) presented at the center of a black background. The two prototypes were defined by opposite directions of coherent motion: upward (90°) and downward (270°), each at 100% coherence. During practice sessions, all participants achieved 100% accuracy in categorizing the prototypical stimuli.

##### Global motion continuum

Nine intermediate global motion stimuli with 10% increments were generated by parametrically varying the relative proportion of dots moving downward (from 10% downward to 90 % downward). In each stimulus, dots moved toward one direction (e.g., downwards) at full coherence (coherence = 1.0), such that perceptual ambiguity arose from the relative proportion of the dots rather than within-population directional noise.

### 5.3 Procedure

#### Task Overview

Participants performed a series of two-alternative forced-choice (2AFC) categorization tasks across four domains: identity, emotion, speech, and global motion. The trial structure remained constant across domains, the stimulus modality and number of sessions were different between two groups.

Deaf participants completed four categorization tasks (seven sessions in total) on the visual modality: i. identity task including dynamic face and static face sessions separately, ii, expression task involving dynamic face and static face sessions separately, iii, speech task including normal face and reversed face sessions separately, and iv, global motion task.

Hearing participants completed four categorization tasks (thirteen sessions in total) across visual, auditory, and audiovisual modalities: i. identity task including facial-dynamic, facial-static, vocal, facial-vocal sessions separately, ii, expression task involving facial-dynamic, facial-static, vocal, facial-vocal sessions separately, iii, speech task including facial-normal, facial-reversed, vocal, facial-vocal sessions separately, and iv, global motion task.

#### Trial structure and task

In all the session, each trial began with a central fixation cross (0.5 s), followed by the presentation of the stimulus (2∼3s), and then a two-alternative forced choice screen (2s).

Participants were instructed to categorize the stimulus by pressing the “F” or “J” keys using the left and right index fingers. Response mappings were counterbalanced within task and within participant to control for motor bias. Each session consisted of 270 trials, with each of the nine stimulus intensity levels repeated 30 times in a pseudo-randomized order.

All participants completed a practice session using the prototypical stimuli prior to the main experiment. A performance criterion of 100% accuracy was required to proceed. The average duration was around 40 minutes for each facial task, 20 minutes for the global motion task, and 20 minutes for each audiovisual (or auditory) task. Deaf participants completed all the tasks in two days (around 160 minutes in total), hearing participants (around 280 minutes in total) in three days. The numbers of participants completing each task were provided in table S7.

### 5.4 Data analysis

#### Data preprocessing and exclusion criteria

Before the formal data analysis, trials with no response were excluded from the dataset (accounting for approximately 0.79% of total trials). Participants were excluded if their categorization accuracy for the prototypical stimuli (the 10% and 90% morphing endpoints) fell below the group mean by more than three standard deviations (mean ± 3 SD). This procedure ensured that all participants included in the final analysis could reliably categorize prototypical categories before their psychometric parameters were estimated.

For each participant and session, we calculated the mean accuracy (e.g., the proportions of choosing happy [or angry] for 90% happy [or 90% angry] face in the expression categorization) on the two prototypical stimuli, and the proportions of the target choice option (e.g., the proportions of choosing ‘happy’ in the expression categorization) at each of the nine stimuli intensities.

#### Psychometric Curve Fitting

Response proportions for the target choice option were fitted with cumulative Gaussian function using the Palamedes toolbox (version 1.11.11) in MATLAB.

The psychometric function is defined as:

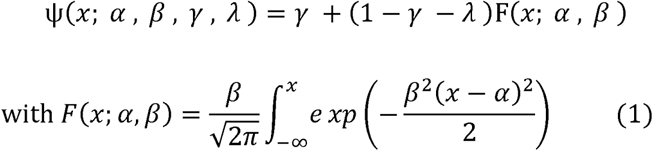

In the function, α is the threshold (point of subjective equality, PSE), reflecting categorical decision boundaries. β is the slope indexing perceptual sensitivity to graded variations in stimulus intensity. γ and λ are the guessing rate and lapse rate to account for non-sensory errors. Ou analysis focused on the slope and threshold parameters. In addition, participants were excluded from the statistical analysis if i. the psychometric function failed to converge during the fitting procedure or the goodness-of-fit was poor, defined as a *p* < 0.05 based on 1,000 bootstrap simulations.

#### Statistical Comparison

To quantify the contributions of facial and vocal signals and their integration, we analyzed data from the hearing group using Linear Mixed-Effects (LME) models in R (R version 4.4.2) with the lme4 and emmeans packages. Separate models were constructed for identity, expression, and speech tasks. In each model, the dependent variable was either the response accuracy, the threshold parameter, or the slope parameter. The fixed factor was stimulus modality (facial, vocal, and facial-vocal), and the random variable was participant. These analyses were restricted to data from participants who completed the task in all three modalities. The model is expressed as:

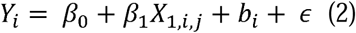

- *Y_i_*: the response accuracy, the threshold parameter, or the slope parameter for participant *i*
- β*_0_,* β*_1_:* fixed-effect coefficients for the intercept and modality
- *b_i_* ∼ *N (0,* σ*^2^)*: random intercept accounting for inter-individual variability
-_: the residual error

To evaluate whether auditory deprivation selectively alters perceptual sensitivity or categorical boundaries, we performed group-level LME analyses for the three facial tasks (identity, expression, and speech) with separate models for each task.

In each model, the dependent variable was either the response accuracy, the threshold parameter, or the slope parameter. The fixed factor was group (deaf group and hearing group) and stimulus type (dynamic vs. static in the facial identity and expression, and normal vs. reversed for speech), and the random variable was participant. The model is expressed as:

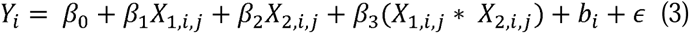

The definitions of parameters were the same as equation (2). Significance levels for these multiple comparisons were adjusted using the Bonferroni correction with a family-wise error rate of α = 0.05.

For the global motion task, group differences were assessed using independent samples t-tests. These analyses only included participants who completed both sessions (e.g., dynamic and static sessions) within a specific task (e.g., identity).

To explore the relationship between perceptual performance and demographic or cognitive factors, we fitted generalized linear models predicting sensitivity (i.e., slope) from age and Raven’s Standard Progressive Matrices score in both groups. Within the deaf group, additional predictors included age of hearing loss onset, sign-language proficiency, and lip-reading proficiency.

#### Contral analysis

To assess the robustness of our findings, we conducted Four control analyses. First, we examined cross-task consistency of the perceptual categorization (i.e., slope and threshold parameters) using *Spearman’s r* coefficient. Then, we repeated the LME model analysis and T test with slope parameters as the dependent variable, group and stimulus type as fixed factors, participants as random variable, and age and Raven’s scores as covariates for each task. After that, we performed LME analysis using data from participants who completed all dynamic visual sessions. In the LME model, the slope parameterwas the dependent variable, stimulus type (dynamic facial identity, dynamic facial expression, normal facial speech, dots-cloud) and group (deaf group and hearing group) were fixed factors, and participant was the random variable. Finally, we repeated the LME analysis and T tests on slope and threshold parameters and cross□task correlations analyses in a subset of deaf and hearing participants matched for age and Raven’s scores to control for potential sampling bias.

