## Supplementary materials for "Domain-Specific Functional Plasticity of Visual Processing Constrained by General Cognitive Ability in Deaf Individuals"

### 1. Supplementary results

#### 1.1 Facial-vocal facilitation in all communicative domains in hearing group

##### 1.1.1 Identity perception

***Response accuracy***. Within the hearing group (table S1), accuracy for prototypical facial-vocal identity stimuli (98.9% ± 1.8%) was comparable to that for facial stimuli (98.4% ± 2.3%), and significantly higher than accuracy on vocal stimuli (98% ± 2.7%; $t_{(198)} = 2.90$, $p = 0.01$, CI $_{95\%} =$ [0.1%, 1.6%]). The difference between facial and vocal accuracy did not reach significance.

***Slope parameter.*** Slopes for facial-vocal identity stimuli (1.58±0.47) were significantly steeper than those for facial stimuli (1.24±0.46; $t_{(146)} = 5.58$, $p_{Bonferroni} = 3.43 \times 10^{-7}$, CI $_{95\%} =$ [0.20, 0.49]; Figure S3A), which in turn were significantly steeper than the slopes for vocal stimuli (1±0.36; $t_{(146)} = 3.77$, $p_{Bonferroni} = 0.0007$, CI $_{95\%}$ = [0.08, 0.28]).

***Threshold parameter.*** Thresholds for facial-vocal identity stimuli (4.48±0.56) were significantly lower than those for facial stimuli (4.79±0.7; $t_{(146)} = 3.15$, $p_{Bonferroni} = 0.006$, CI $_{95\%} =$ [0.07, 0.54]; Figure S3A) and were comparable to those for vocal stimuli (4.69±0.71). The difference between facial and vocal thresholds did not reach significance.

##### 1.1.2 Emotion perception

***Response accuracy***. Within the hearing group (Table S1), accuracy for prototypical facial-vocal emotion (99.1% ± 1.8%; $t_{(252)} = 4.84$, $p_{Bonferroni} = 6.75 \times 10^{-6}$, CI $_{95\%} =$ [0.5%, 1.5%]) and for facial emotion (99.3% ± 1.2%; $t_{(252)} = 5.98$, $p_{Bonferroni} = 2.34 \times 10^{-8}$, CI $_{95\%} =$ [0.8%, 1.8%]) were higher than accuracy for vocal emotion (98.1% ± 2.4%). No significant difference was observed between facial-vocal and facial accuracy.

***Slope parameter***. Slopes for facial-vocal emotion (1.65±0.50) were significantly steeper than those for facial emotion (1.3±0.41; $t_{(208)} = 6.96$, $p_{Bonferroni} = 1.32 \times 10^{-10}$, CI $_{95\%} =$ [0.23, 0.47]; Figure S3B), which in turn were significantly steeper than slopes for vocal emotion (0.97±0.33; $t_{(208)} = 6.73$, $p_{Bonferroni} = 4.74 \times 10^{-10}$, CI $_{95\%}$ = [0.21, 0.45]).

***Threshold parameter.*** Thresholds for facial-vocal emotion (5.0±0.61; $t_{(208)} = -3.12$, $p_{Bonferroni} = 0.006$, CI $_{95\%} =$ [-0.35, -0.05]; Figure S3B) and facial emotion (5.05±0.64; $t_{(208)} = -2.43$, $p_{Bonferroni} = 0.048$, CI $_{95\%} =$ [-0.31, -0.001]) were significantly lower than vocal emotion (5.2±0.64).

#### 1.1.3 Speech perception

***Response accuracy***. Within the hearing group (Table S1), categorization accuracy for facial-vocal speech (99.6% ± 1.1%; $t_{(210)}$ = -8.49, $p_{Bonferroni}$ = 1.15 × $10^{-14}$, CI $_{95\%=}$ [2.2%, 3.9%]) and vocal speech (99. 4%± 1.3%; $t_{(210)}$ = -8.12, $p_{Bonferroni}$ = 1.16 × $10^{-13}$, CI $_{95\%=}$ [2.1%, 3.8%])) was significantly higher than accuracy for normal facial articulation (96.5% ±4.5%). No significant difference was observed between facial-vocal and vocal speech.

***Slope parameter.*** Slopes for facial-vocal speech stimuli (1.77±0.6; $t_{(82)}$ = 6.41, $p_{Bonferroni}$ = 2.65 × $10^{-8}$, CI $_{95\%=}$ [0.47, 1.04]; Figure 3C) and vocal speech(1.63±0.68; $t_{(82)}$ = 5.2, $p_{Bonferroni}$ = 4.39 × $10^{-6}$, CI $_{95\%=}$ [0.32, 0.9]) were significantly steeper than those for facial stimuli (1.02±0.33). No significant difference was observed between facial-vocal speech and vocal speech. These results align with prior findings that vocal cues provide more reliable speech information than facial signals.

***Threshold parameter.*** Thresholds for facial-vocal speech (5.46±0.56) were significantly higher than those for facial speech (5.02±0.63; $t_{(82)}$ = 3.62, $p_{Bonferroni}$ = 0.001, $CI_{95\%}$ = [0.14, 0.74]; Figure 3C). No significant difference was observed between facial-vocal and vocal speech (5.28±0.63).

#### 1.2 Individual differences in perceptual threshold. predicted by general cognitive ability

Raven's score significantly predicted threshold parameters for static ($\beta$ = 0.003, $t$ = 2.26, $p$ = 0.03) and dynamic ($\beta$ = 0.003, $t$ = 2.41, $p$ = 0.02) facial expression categorization. Age of hearing loss significantly predicted the threshold for static facial expressions ($\beta$ = 0.02, $t$ = 2.38, $p$ = 0.02). Lip-reading proficiency predicted the threshold for static facial expressions ($\beta$ = -0.04, $t$ = 2.04, $p$ = 0.04).

### 2. Supplementary figures

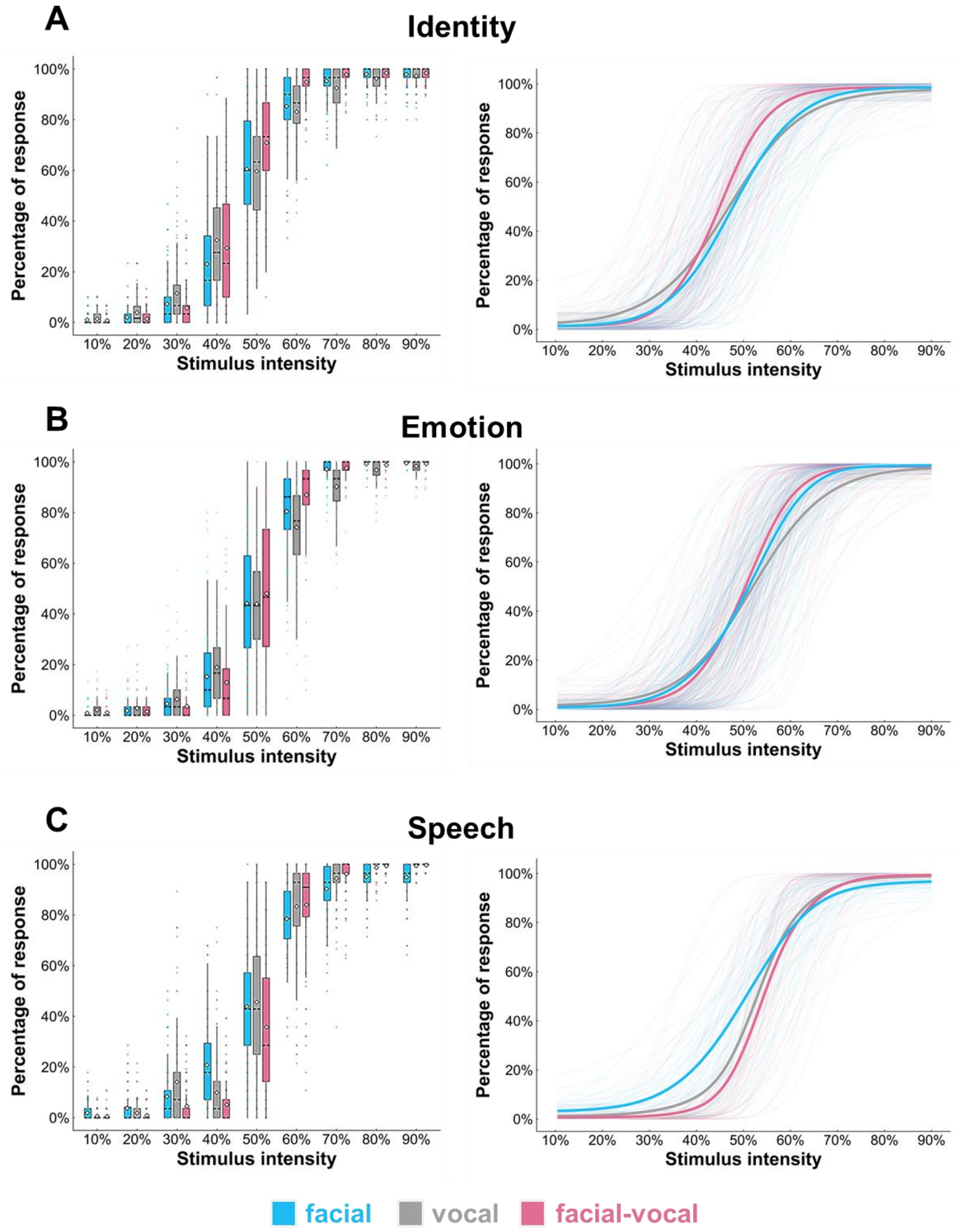


**Figure S1. Response proportions (left) and psychometric curves (right) in the hearing group.** (A) Proportions of choosing "female" in the dynamic facial (blue), vocal (gray), and facial-vocal (red) identity sessions. Thick curves represent the group mean, and thin curves represent individual participants. (B) Proportions of choosing "happy" in the dynamic facial, vocal, and facial-vocal emotion sessions. (C) Proportions of choosing "da" in the static facial, vocal, and facial-vocal speech sessions.

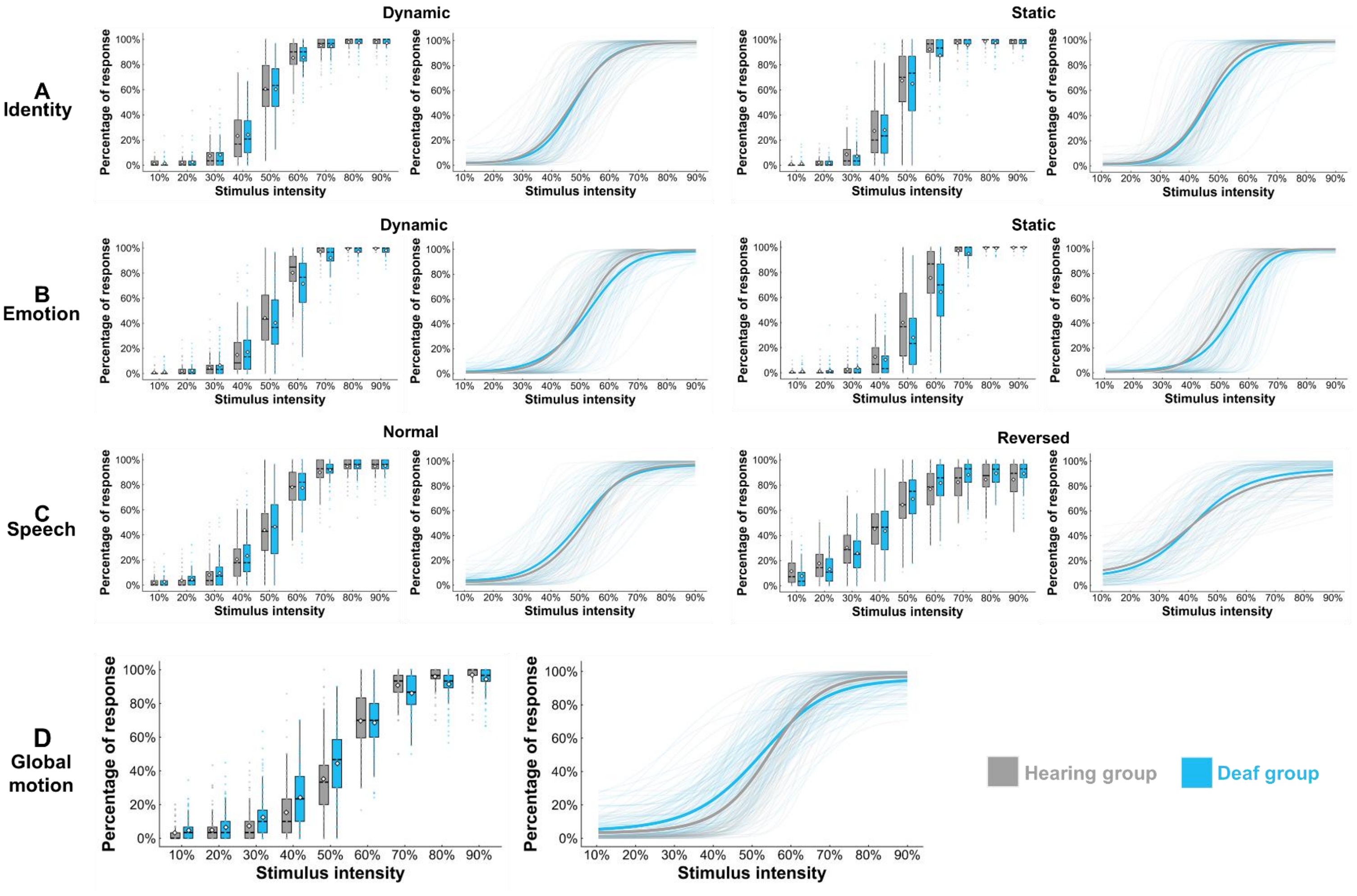


**Figure S2. Response proportions (left) and psychometric curves (right) across visual tasks in hearing (gray) and deaf (blue) groups.** (A) Proportions of choosing "female" in the dynamic and static facial identity task. Thick curves represent the group mean, and thin curves represent individual participants. (B) Proportions of choosing "happy" in dynamic and static facial expression task. (C) Proportions of choosing "da" in normal and reversed facial speech task. (D) Proportions of choosing "downward" in the global motion task.

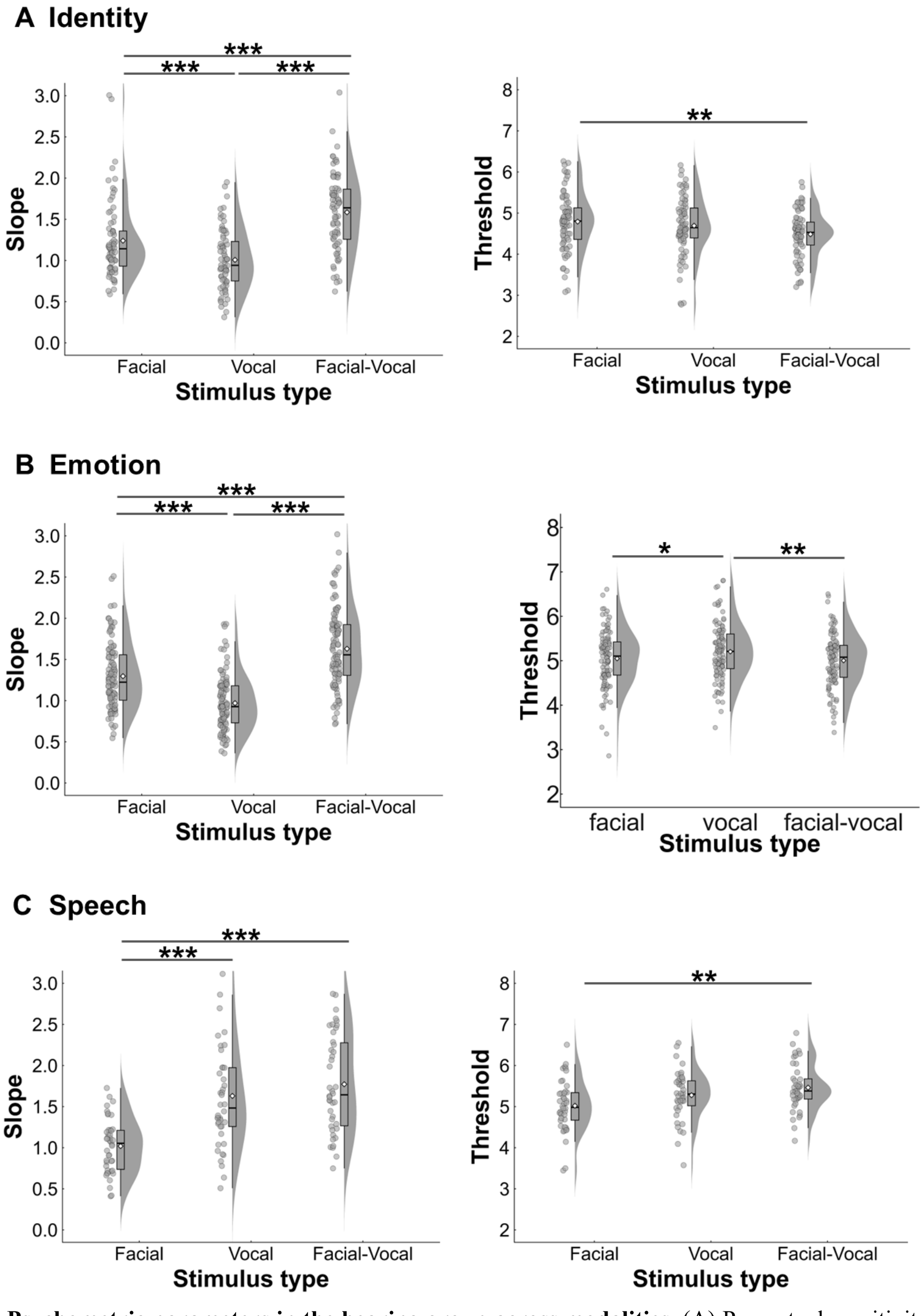


**Figure S3. Psychometric parameters in the hearing group across modalities**. (A) Perceptual sensitivity (indexed by slope) and categorization boundaries (indexed by threshold) for facial, vocal, and facial-vocal identity. (B) Perceptual sensitivity and categorization boundaries for facial, vocal, and facial-vocal emotion. (C) Perceptual sensitivity and categorization boundaries for facial, vocal, and facial-vocal speech. In all domains, audiovisual integration enhanced perceptual sensitivity relative to unisensory conditions.

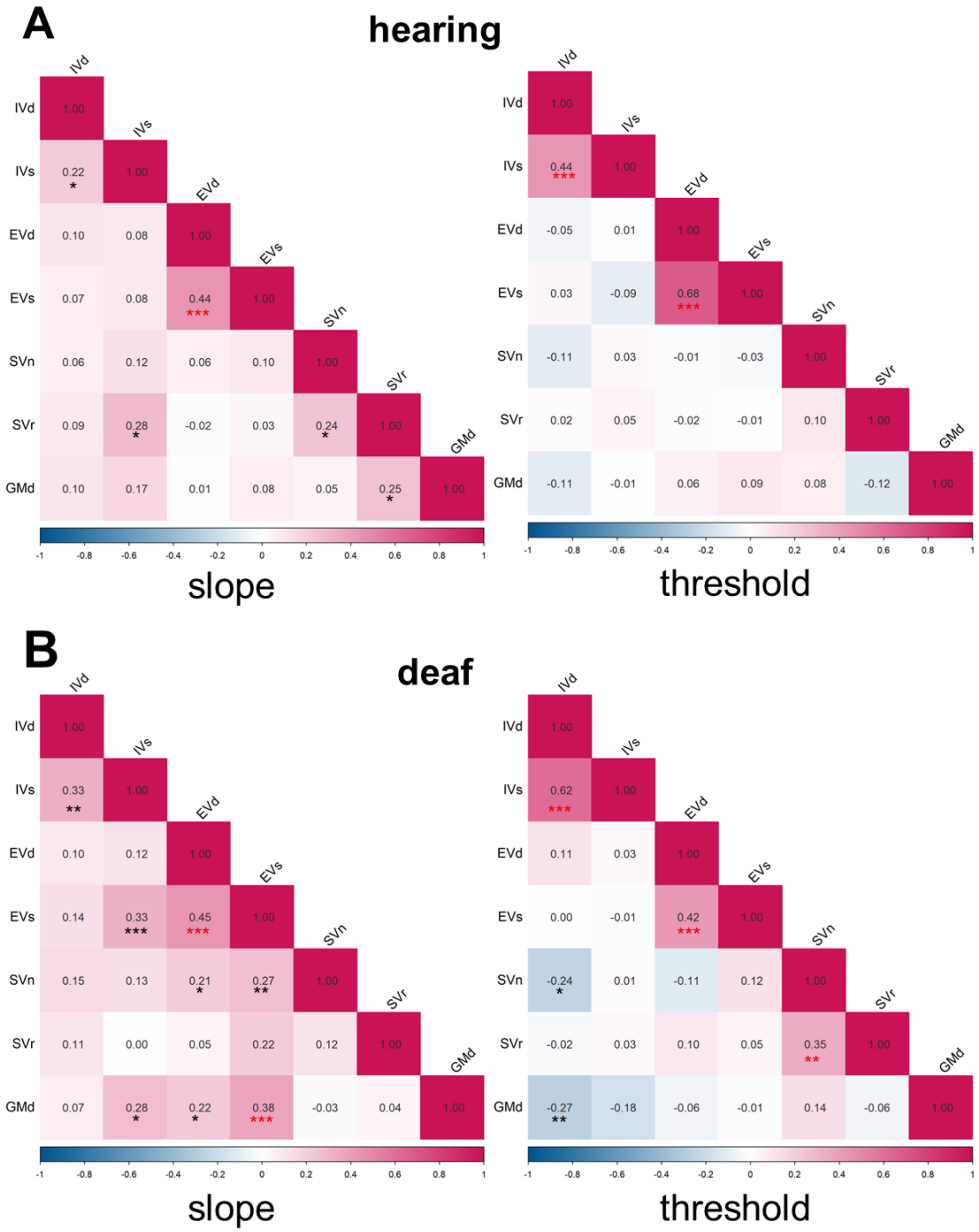


**Figure S4. Pairwise correlations of perceptual sensitivity (indexed by slope) and categorization boundaries (indexed by threshold) across all visual tasks.** (A) Deaf group. (B) Hearing group. Black asterisks indicate significant correlations, and red asterisks indicate correlations that remained significant after Bonferroni correction ($\alpha$ = 0.05). Abbreviations: IVd = identity task with dynamic visual stimuli; IVs = identity task with static visual stimuli; EVd = emotion task with dynamic visual stimuli; EVs = emotion with static visual stimuli; SVn = speech task with normal visual stimuli; SVr = speech task with reversed visual stimuli; GMd = global motion with dynamic stimuli. *, $p$ <0.05; **, $p$ <0.01, ***, $p$ <0. 001.

### 3. Supplementary tables

**Table S1. Response proportions across all stimulus intensities in facial, vocal, and facial-vocal sessions in the hearing group**

| task | session | group | N | intensity of stimulus 10% | 20% | 30% | 40% | 50% | 60% | 70% | 80% | 90% | ACC |
|---|---|---|---|---|---|---|---|---|---|---|---|---|---|
| **identity** | **facial** | hearing | 100 | 1.0±2.1 | 2.1±3.5 | 7.3±9.2 | 23.1±20.8 | 60.5±25.5 | 85.3±15.0 | 95.3±6.9 | 98.0±3.7 | 97.9±3.9 | **98.4±2.3** |
| | **vocal** | | | 1.4±2.6 | 3.9±5.8 | 11.5±14.6 | 32.5±21.7 | 59.7±20.0 | 83.1±13.0 | 92.5±8.2 | 96.3±5.3 | 97.3±3.9 | **98.0±2.7** |
| | **facial-vocal** | | | 0.8±1.7 | 1.4±2.6 | 5.7±9.0 | 29.4±23.8 | 70.9±22.1 | 94.8±7.7 | 97.7±3.8 | 98.5±3.2 | 98.5±2.7 | **98.9±1.8** |
| **emotion** | **facial** | hearing | 127 | 0.8±2.1 | 1.8±3.9 | 4.6±6.9 | 15.3±17.1 | 44.2±24.7 | 80.4±17.7 | 97.1±4.6 | 99.2±1.9 | 99.5±1.2 | **99.3±1.2** |
| | **vocal** | | | 2.1±3.4 | 2.8±4.9 | 6.3±8.8 | 18.9±14.5 | 43.9±20.7 | 74.1±17.5 | 90.2±10.3 | 96.8±4.9 | 98.3±2.8 | **98.1±2.4** |
| | **facial-vocal** | | | 1.0±2.3 | 1.4±2.6 | 3.5±6.1 | 13.0±16.2 | 48.0±28.2 | 87.0±16.6 | 97.5±4.1 | 98.8±2.5 | 99.2±2.0 | **99.1±1.8** |
| **speech** | **facial** | hearing | 106 | 1.9±3.4 | 3.8±6.1 | 8.3±10.6 | 20.8±16.8 | 43.9±21.7 | 78.5±14.4 | 90.3±10.6 | 95.1±6.6 | 94.9±7.5 | **96.5±4.5** |
| | **vocal** | | | 0.6±1.7 | 1.8±3.8 | 14.1±19.3 | 9.9±14.7 | 45.6±26.2 | 83.3±19.8 | 94.4±9.4 | 98.7±2.9 | 99.4±1.6 | **99.4±1.3** |
| | **facial-vocal** | | | 0.5±1.8 | 0.9±2.2 | 4.5±7.9 | 5.1±8.7 | 35.7±26.8 | 84.0±19.1 | 96.2±7.4 | 99.4±1.6 | 99.7±1.0 | **99.6±1.1** |

Note: Because not all participants completed every task, the sample size varied across tasks. For each task, we included only the participants who completed all sessions (i.e., facial, vocal, and facial-vocal).

**Table S2. Psychometric parameters for facial, vocal, and facial–vocal sessions in the hearing group**

| task | session | group | N | slope | threshold |
|---|---|---|---|---|---|
| **identity** | **facial** | hearing | 74 | 1.24 ± 0.46 | 4.79±0.7 |
| | **vocal** | | | 1.01 ± 0.36 | 4.69±0.71 |
| | **facial-vocal** | | | 1.58 ± 0.47 | 4.48±0.56 |
| **emotion** | **facial** | hearing | 103 | 1.30 ± 0.41 | 5.05±0.64 |
| | **vocal** | | | 0.97 ± 0.33 | 5.20±0.64 |
| | **facial-vocal** | | | 1.65 ± 0.50 | 5.00±0.61 |
| **speech** | **facial** | hearing | 42 | 1.02 ± 0.33 | 5.02±0.63 |
| | **vocal** | | | 1.63 ± 0.68 | 5.28±0.6 |
| | **facial-vocal** | | | 1.77 ± 0.60 | 5.46±0.56 |

Note: For each task, we included only the participants who completed all sessions (i.e., facial, vocal, and facial-vocal), whose psychometric functions successfully converged, and whose goodness-of-fit was acceptable ($p > 0.05$).

**Table S3. Response proportions across all stimulus intensities in facial sessions across all perceptual domains in deaf and hearing groups**

| task | stimulus | group | N | intensity of stimulus 10% | 20% | 30% | 40% | 50% | 60% | 70% | 80% | 90% | ACC |
|---|---|---|---|---|---|---|---|---|---|---|---|---|---|
| identity | dynamic | hearing | 106 | 1.2±2.2 | 2.1±3.5 | 7.5±9.1 | 23.2±20.6 | 60.5±25.3 | 85.2±15.3 | 95.0±7.2 | 97.9± 3.8 | 97.9±3.8 | **98.4±2.4** |
| | | deaf | 92 | 1.4±3.4 | 2.3±5.9 | 8.2±10.9 | 24.1±17.5 | 60.3±20.5 | 85.5±13.2 | 94.4±7.2 | 97.3±5.3 | 97.0±6.9 | **97.8±4.5** |
| | static | hearing | 106 | 0.9±1.9 | 2.0±3.5 | 8.6±11.0 | 27.2±23.5 | 67.4±23.6 | 92.2±11.1 | 97.3±4.2 | 98.6± 3.1 | 98.2±2.8 | **98.7±1.8** |
| | | deaf | 92 | 1.2±2.9 | 1.6±2.8 | 7.1±11.3 | 27.9±23.6 | 64.8±27.2 | 87.4±17.4 | 95.6±8.9 | 97.5± 5.0 | 98.2±3.4 | **98.5±2.6** |
| emotion | dynamic | hearing | 122 | 0.8±2.1 | 2.0±4.2 | 4.6±7.0 | 14.9±16.7 | 44.3±25.0 | 80.3±17.7 | 96.9±5.7 | 99.3±1.9 | 99.5±1.2 | **99.3± 1.2** |
| | | deaf | 123 | 1.1±2.9 | 2.3±4.1 | 6.6±10.1 | 17.1±17.8 | 40.3±25.1 | 71.5±22.1 | 92.2±10.0 | 97.2±5.2 | 98.1±3.3 | **98.5±2.5** |
| | static | hearing | 122 | 0.7±1.7 | 1.2±2.9 | 2.8±5.2 | 12.6±15.4 | 39.9±29.5 | 75.5±27.0 | 97.4±7.9 | 99.3±1.5 | 99.6±1.2 | **99.4±1.1** |
| | | deaf | 123 | 1.1±2.4 | 1.6±4.3 | 3.8±8.2 | 10.4±15.1 | 28.0±25.1 | 64.4±27.8 | 94.9±9.5 | 99.2±1.8 | 99.5±1.2 | **99.2±1.4** |
| speech | normal | hearing | 108 | 1.9±3.3 | 3.6±6.0 | 8.1±10.6 | 20.5±16.5 | 43.8±22.2 | 78.1±15.2 | 89.9±11.2 | 94.7±7.1 | 94.7±7.5 | **96.4±4.6** |
| | | deaf | 93 | 2.6±4.4 | 4.5±6.6 | 9.5±11.4 | 23.4±20.4 | 46.5±25.5 | 77.6±18.8 | 91.8± 9.3 | 94.7±7.5 | 95.1±6.3 | **96.3±4.5** |
| | reverse | hearing | 108 | 11.5±12.1 | 17.7±14.0 | 30.2±17.5 | 45.1±20.0 | 64.3±20.9 | 76.8±17.5 | 82.2±15.3 | 84.4±14.5 | 84.3±14.8 | **86.4±11.5** |
| | | deaf | 93 | 7.7±9.1 | 13.1±11.7 | 25.7±15.3 | 43.4±19.8 | 68.9±19.5 | 81.6±16.9 | 88.1±12.2 | 89.3±9.3 | 89.1±11.6 | **90.7±8.7** |
| global motion | dynamic | hearing | 123 | 3.2±4.8 | 4.9±7.7 | 7.3±9.8 | 15.3±16.3 | 35.2±20.6 | 69.6±17.0 | 90.8±9.6 | 96.0±6.0 | 96.9±5.3 | **96.8±4.3** |
| | | deaf | 117 | 4.7±6.4 | 6.5±8.8 | 12.5±12.3 | 24.2±16.7 | 44.4±20.2 | 68.5±16.0 | 86.0±10.6 | 91.4±9.1 | 94.4±7.5 | **94.8±6.2** |

Note: Because not all participants completed every task, the sample size varied across tasks. For each task, we included only the participants who completed all sessions (i.e., facial, vocal, and facial-vocal).

**Table S4. Psychometric parameters of facial sessions in deaf and hearing groups**

| task | stimulus | group | N | slope | threshold |
|---|---|---|---|---|---|
| **identity** | **dynamic** | hearing | 89 | 1.24±0.47 | 4.74±0.71 |
| | | deaf | 80 | 1.21±0.44 | 4.79±0.56 |
| | **static** | hearing | 89 | 1.41±0.57 | 4.53±0.61 |
| | | deaf | 80 | 1.42±0.51 | 4.72±0.79 |
| **expression** | **dynamic** | hearing | 100 | 1.30±0.40 | 5.05±0.64 |
| | | deaf | 101 | 1.11±0.40 | 5.22±0.78 |
| | **static** | hearing | 100 | 1.60±0.54 | 5.16±0.67 |
| | | deaf | 101 | 1.53±0.62 | 5.43±0.76 |
| **speech** | **normal** | hearing | 80 | 1.06±0.43 | 5.12±0.66 |
| | | deaf | 78 | 1.04±0.38 | 4.96±0.89 |
| | **reverse** | hearing | 80 | 0.75±0.33 | 4.33±1.23 |
| | | deaf | 78 | 0.82±0.33 | 4.28±0.95 |
| **global motion** | **dynamic** | hearing | 110 | 1.11±0.38 | 5.37±0.61 |
| | | deaf | 111 | 0.86±0.36 | 5.15±0.73 |

Note: For each task, we included only the participants who completed all sessions (i.e., facial, vocal, and facial-vocal), whose psychometric functions successfully converged, and whose goodness-of-fit was acceptable ($p > 0.05$).

**Table S5. Demographics of the deaf and hearing participants with matched age and RSPM Score**

| | **Hearing controls (*N* = 33)** | **Deaf participants (*N* =27)** | **Statistics** | ***p*-value** |
|---|---|---|---|---|
| **Gender: males/females (n)** | 13/ 20 | 18 / 9 | / | / |
| **Age (years)** | 21.45 ± 2.15 | 22.48 ± 2.155 | 0.84 | 0.07 |
| **RSPM Score** | 106.14 ± 10.593 | 100.65 ± 11.823 | 11.9 | 0.06 |
| **Handedness: right/left (n)** | 31/2 | 27/0 | / | / |
| **Age of HL onset (years)** | / | 1.24 ± 1.74 | / | / |
| **Severe-to-profound HL (%)** | / | 100% | / | / |
| **Sign language users (%)** | / | 92.6 | / | / |
| **Age of SL onset (years)** | / | 8.64 ± 3.65 | / | / |
| **Proficiency of SL (1-10)** | / | 6.15 ± 2.44 | / | / |
| **Proficiency of lip-reading (1-5)** | / | 2.78 ± 0.97 | / | / |

RSPM = Raven’s Standard Progressive Matrices; HL = hearing loss; SL = sign language; Self-reported SL proficiency: 1 = no ability, 10 = native/near-native fluency; Self-reported lip-reading proficiency: 1 = cannot understand any conversations, 5 = can understand most conversations; “/” not applicable to this group.

**Table S6. Detailed demographic information for all participants**

| Number | ID | group | age | gender | Raven's Score | handedness | education level | age of hearing loss | degree of hearing loss | hearing aid usage | sign language proficiency | lip-reading proficiency | cause of hearing loss |
|---|---|---|---|---|---|---|---|---|---|---|---|---|---|
| 1 | D001 | deaf | 21 | female | 102.36 | right | college | 0 | severe | everyday | 7 | 4 | ototoxicity |
| 2 | D002 | deaf | 20 | female | 124.67 | right | college | 2 | severe | everyday | 7 | 3 | sudden |
| 3 | D003 | deaf | 22 | female | 95.22 | right | college | 3 | profound | never | 5 | 2 | unclear |
| 4 | D004 | deaf | 22 | female | 93.54 | right | college | 1 | profound | never | 5 | 2 | unclear |
| 5 | D005 | deaf | 21 | male | 96.84 | right | college | 1 | severe | never | 8 | 3 | unclear |
| 6 | D006 | deaf | 22 | female | 87.38 | right | college | 1 | profound | never | 6 | 2 | unclear |
| 7 | D007 | deaf | 32 | female | 102.36 | right | college | 1 | severe | never | 8 | 2 | unclear |
| 8 | D008 | deaf | 20 | female | 95.22 | right | college | 0 | severe | never | 6 | 2 | ototoxicity |
| 9 | D009 | deaf | 24 | male | 88.67 | right | college | 0 | severe | never | 5 | 2 | unclear |
| 10 | D010 | deaf | 18 | female | 86.59 | right | college | 1 | profound | never | 6 | 3 | unclear |
| 11 | D011 | deaf | 27 | male | 98.43 | right | college | 3 | severe | sometimes | 6 | 2 | sudden |
| 12 | D012 | deaf | 22 | male | 100 | right | college | 1 | profound | never | 5 | 2 | unclear |
| 13 | D013 | deaf | 24 | male | 104.78 | left | college | 7 | severe | never | 8 | 3 | sudden |
| 14 | D014 | deaf | 23 | male | 79.89 | right | college | 3 | profound | never | 9 | 1 | sudden |
| 15 | D015 | deaf | 25 | male | / | right | primary | 3 | profound | never | 3 | 1 | sudden |
| 16 | D016 | deaf | 20 | male | 95.22 | right | college | 1 | profound | never | 6 | 2 | sudden |
| 17 | D017 | deaf | 22 | male | 93.54 | right | college | 2 | profound | never | 5 | 2 | unclear |
| 18 | D018 | deaf | 23 | male | 95.22 | right | college | 0 | profound | never | 7 | 1 | unclear |
| 19 | D019 | deaf | 21 | female | 91.77 | right | college | 1 | severe | everyday | 6 | 3 | sudden |
| 20 | D020 | deaf | 19 | female | 124.67 | right | college | 4 | severe | everyday | 5 | 3 | unclear |
| 21 | D021 | deaf | 21 | female | 80.78 | right | college | 0 | profound | never | 7 | 3 | ototoxicity |
| 22 | D022 | deaf | 21 | female | 76.68 | right | college | 1 | profound | everyday | 5 | 2 | unclear |
| 23 | D023 | deaf | 29 | female | 100 | right | college | 0 | severe | never | 6 | 2 | sudden |
| 24 | D024 | deaf | 25 | male | 104.78 | right | college | 1 | severe | never | 6 | 2 | heredity |
| 25 | D026 | deaf | 27 | female | 100 | right | college | 0 | severe | everyday | 7 | 3 | unclear |
| 26 | D027 | deaf | 29 | female | 98.43 | right | college | 2 | profound | never | 6 | 2 | heredity |
| 27 | D028 | deaf | 32 | female | 110.12 | right | high | 1 | profound | never | 6 | 3 | heredity |
| 28 | D029 | deaf | 27 | male | 100 | left | college | 1 | profound | never | 6 | 3 | heredity |
| 29 | D031 | deaf | 29 | female | 89.88 | right | college | 1 | profound | never | 6 | 1 | ototoxicity |

| | | | | | | | | | | | | | |
|---|---|---|---|---|---|---|---|---|---|---|---|---|---|
| 30 | D032 | deaf | 27 | female | 102 | right | college | 0 | severe | everyday | 7 | 3 | heredity |
| 31 | D034 | deaf | 25 | male | / | right | college | 3 | profound | never | 6 | 3 | ototoxicity |
| 32 | D035 | deaf | 26 | female | 100 | right | college | 2 | severe | everyday | 10 | 3 | unclear |
| 33 | D036 | deaf | 22 | male | 78.92 | right | college | 0 | severe | everyday | 6 | 3 | sudden |
| 34 | D037 | deaf | 21 | male | 93.54 | right | college | 1 | severe | everyday | 8 | 4 | unclear |
| 35 | D038 | deaf | 30 | male | 120 | right | college | 0 | severe | never | 5 | 3 | heredity |
| 36 | D039 | deaf | 25 | female | 95.22 | right | graduate | 0 | severe | sometimes | 9 | 4 | heredity |
| 37 | D040 | deaf | 24 | female | 120 | right | college | 0 | severe | everyday | 9 | 4 | heredity |
| 38 | D041 | deaf | 19 | male | 100 | right | college | 1.5 | severe | never | 8 | 2 | heredity |
| 39 | D042 | deaf | 25 | female | 102.36 | right | college | 0 | profound | never | 8 | 1 | ototoxicity |
| 40 | D043 | deaf | 26 | male | 89.88 | right | college | 0 | severe | never | 6 | 3 | unclear |
| 41 | D044 | deaf | 26 | male | 93.54 | right | high | 3 | profound | sometimes | 8 | 3 | ototoxicity |
| 42 | D045 | deaf | 37 | male | 93.78 | right | primary | 0 | profound | never | 5 | 1 | sudden |
| 43 | D046 | deaf | 19 | male | 87.75 | right | college | 0 | profound | never | 7 | 3 | unclear |
| 44 | D047 | deaf | 27 | female | 95.22 | right | college | 2 | profound | everyday | 5 | 3 | sudden |
| 45 | D048 | deaf | 30 | male | 100 | right | high | 3 | severe | never | 8 | 3 | ototoxicity |
| 46 | D049 | deaf | 31 | male | 97.3 | right | high | 0 | profound | never | 5 | 3 | unclear |
| 47 | D050 | deaf | 21 | male | 85.98 | right | college | 0 | profound | never | 8 | 3 | unclear |
| 48 | D051 | deaf | 20 | female | 96.84 | right | college | 0 | profound | never | 5 | 1 | unclear |
| 49 | D052 | deaf | 30 | female | 96.84 | right | college | 0 | profound | sometimes | 8 | 3 | heredity |
| 50 | D053 | deaf | 24 | male | 120 | right | college | 1 | severe | sometimes | 5 | 3 | heredity |
| 51 | D054 | deaf | 26 | female | 88.67 | right | college | 3 | severe | everyday | 10 | 4 | ototoxicity |
| 52 | D055 | deaf | 25 | female | 77.86 | right | college | 2 | severe | never | 6 | 2 | unclear |
| 53 | D056 | deaf | 25 | female | 80.78 | right | college | 7 | profound | never | 7 | 3 | unclear |
| 54 | D057 | deaf | 31 | male | 102.36 | right | college | 2.5 | severe | never | 5 | 2 | sudden |
| 55 | D058 | deaf | 31 | female | 75.33 | right | junior | 6 | profound | sometimes | 6 | 3 | unclear |
| 56 | D059 | deaf | 28 | male | 96.84 | right | college | 3 | profound | everyday | 6 | 3 | sudden |
| 57 | D060 | deaf | 22 | male | 91.77 | right | college | 0 | profound | everyday | 5 | 3 | heredity |
| 58 | D061 | deaf | / | male | / | / | / | / | / | / | / | / | / |
| 59 | D063 | deaf | 28 | male | / | right | college | 0 | profound | never | 2 | 1 | unclear |
| 60 | D064 | deaf | 23 | female | 95.22 | right | college | 0 | profound | never | 7 | 3 | unclear |
| 61 | D065 | deaf | 23 | female | 104.78 | right | college | 0 | severe | never | 7 | 2 | unclear |

| | | | | | | | | | | | | | |
|---|---|---|---|---|---|---|---|---|---|---|---|---|---|
| 62 | D066 | deaf | 24 | female | 76.68 | right | college | 0 | severe | never | 7 | 2 | sudden |
| 63 | D067 | deaf | 28 | female | 114.02 | right | college | 0 | severe | everyday | 4 | 3 | unclear |
| 64 | D068 | deaf | 22 | male | 78.92 | right | college | 0 | profound | never | 6 | 2 | sudden |
| 65 | D069 | deaf | 21 | male | 85.98 | right | college | 0 | profound | never | 5 | 3 | unclear |
| 66 | D070 | deaf | 22 | male | 88.67 | right | college | 3 | profound | never | 8 | 3 | heredity |
| 67 | D071 | deaf | 25 | female | 124.67 | right | college | 0 | profound | everyday | 0 | 4 | sudden |
| 68 | D072 | deaf | 22 | male | 119.22 | right | college | 1 | profound | sometimes | 6 | 3 | sudden |
| 69 | D073 | deaf | 21 | male | 89.88 | right | college | 3 | severe | everyday | 9 | 4 | sudden |
| 70 | D074 | deaf | 24 | female | 95.22 | right | college | 1 | severe | everyday | 0 | 3 | unclear |
| 71 | D075 | deaf | 28 | female | 95.22 | right | high | 1 | severe | everyday | 0 | 3 | unclear |
| 72 | D076 | deaf | 27 | female | 96.84 | right | college | 4 | severe | everyday | 3 | 3 | sudden |
| 73 | D077 | deaf | 28 | male | 95.22 | right | college | 0 | severe | everyday | 8 | 4 | ototoxicity |
| 74 | D078 | deaf | 28 | female | 107.33 | right | college | 0.16 | severe | sometimes | 6 | 3 | unclear |
| 75 | D079 | deaf | 29 | female | 75.33 | right | college | 11 | severe | everyday | 7 | 3 | unclear |
| 76 | D080 | deaf | 26 | female | 104.78 | right | college | 2.5 | severe | never | 6 | 3 | ototoxicity |
| 77 | D081 | deaf | 28 | female | 85.98 | right | college | 0 | severe | sometimes | 4 | 3 | unclear |
| 78 | D082 | deaf | 21 | male | 75.33 | left | college | 3 | profound | never | 5 | 2 | sudden |
| 79 | D083 | deaf | 25 | female | 96.84 | right | college | 0 | profound | never | 8 | 3 | sudden |
| 80 | D084 | deaf | 24 | male | 75.33 | left | high | 0 | severe | never | 6 | 2 | heredity |
| 81 | D085 | deaf | 20 | male | 98.43 | right | high | 0 | profound | never | 7 | 2 | unclear |
| 82 | D086 | deaf | 25 | female | 87.37 | right | college | 0 | profound | never | 5 | 2 | unclear |
| 83 | D087 | deaf | 25 | male | 84.45 | right | high | 1 | profound | never | 5 | 3 | sudden |
| 84 | D088 | deaf | 22 | male | 85.98 | right | college | 1 | profound | sometimes | 10 | 2 | unclear |
| 85 | D089 | deaf | 19 | female | 75.34 | left | college | 1 | profound | everyday | 3 | 4 | sudden |
| 86 | D090 | deaf | 25 | male | 104.8 | right | college | 5 | profound | never | 5 | 1 | sudden |
| 87 | D091 | deaf | 20 | male | 75.33 | right | high | 1 | profound | never | 6 | 2 | sudden |
| 88 | D092 | deaf | 24 | male | 91.77 | right | college | 3 | profound | sometimes | 9 | 1 | sudden |
| 89 | D093 | deaf | 22 | male | 93.54 | right | college | 0 | profound | everyday | 5 | 3 | sudden |
| 90 | D095 | deaf | 27 | male | 75.33 | left | high | 1 | profound | never | 5 | 2 | sudden |
| 91 | D096 | deaf | 23 | male | 95.22 | right | college | 3 | profound | sometimes | 5 | 2 | sudden |
| 92 | D097 | deaf | 22 | male | 95.22 | right | college | 3 | severe | sometimes | 5 | 3 | sudden |
| 93 | D098 | deaf | 24 | female | 95.22 | right | college | 1 | severe | never | 6 | 3 | heredity |

| | | | | | | | | | | | | | |
|---|---|---|---|---|---|---|---|---|---|---|---|---|---|
| 94 | D099 | deaf | 23 | male | 87.38 | right | college | 1 | profound | never | 4 | 1 | sudden |
| 95 | D100 | deaf | 25 | female | 88.67 | right | college | 3 | profound | everyday | 6 | 3 | sudden |
| 96 | D101 | deaf | 20 | male | 76.68 | right | college | 8 | profound | never | 6 | 3 | unclear |
| 97 | D102 | deaf | 26 | male | 114.02 | right | college | 0 | profound | everyday | 7 | 1 | sudden |
| 98 | D103 | deaf | 19 | male | 89.88 | right | high | 3 | profound | never | 3 | 2 | unclear |
| 99 | D104 | deaf | 20 | male | 87.38 | left | college | 0 | profound | everyday | 6 | 4 | sudden |
| 100 | D105 | deaf | 23 | male | 98.43 | right | college | 0 | profound | sometimes | 5 | 3 | unclear |
| 101 | D106 | deaf | 21 | male | 75.33 | right | college | 0 | profound | everyday | 4 | 3 | sudden |
| 102 | D107 | deaf | 25 | female | 91.77 | left | college | 2 | severe | never | 4 | 2 | sudden |
| 103 | D108 | deaf | 25 | male | 124.67 | right | college | 0 | profound | everyday | 6 | 4 | sudden |
| 104 | D109 | deaf | 22 | female | 85.98 | right | college | 2 | profound | never | 4 | 3 | sudden |
| 105 | D110 | deaf | / | male | 110.12 | left | unclear | 1 | profound | never | / | / | unclear |
| 106 | D111 | deaf | 22 | female | 110.11 | right | college | 0 | severe | never | 10 | 2 | unclear |
| 107 | D112 | deaf | 27 | male | 91.77 | right | college | 2 | profound | sometimes | 4 | 1 | sudden |
| 108 | D113 | deaf | 27 | male | / | right | primary | 3 | profound | never | 4 | 3 | sudden |
| 109 | D114 | deaf | 25 | male | / | right | college | 3 | moderate | never | 7 | 3 | sudden |
| 110 | D115 | deaf | 21 | female | 75.33 | left | college | 0 | severe | never | 6 | 2 | sudden |
| 111 | D116 | deaf | 24 | male | 75.33 | left | college | 3 | severe | sometimes | 8 | 5 | sudden |
| 112 | D117 | deaf | 25 | female | 80.78 | left | college | 0 | severe | never | 9 | 3 | unclear |
| 113 | D118 | deaf | 21 | female | 79.89 | right | college | 1 | severe | sometimes | 3 | 3 | sudden |
| 114 | D119 | deaf | 27 | female | 80.78 | right | high | 5 | profound | everyday | 0 | 3 | sudden |
| 115 | D120 | deaf | 19 | female | 75.33 | right | high | 11 | moderate | everyday | 0 | 3 | sudden |
| 116 | D121 | deaf | 25 | male | 119.22 | right | high | 0 | profound | never | 7 | 3 | unclear |
| 117 | D122 | deaf | 23 | male | / | right | high | 8 | profound | never | 10 | 2 | sudden |
| 118 | D123 | deaf | 24 | male | 77.86 | left | junior | 3 | moderate | never | 8 | 3 | unclear |
| 119 | D124 | deaf | 22 | female | 80.78 | left | primary | 2 | severe | never | 8 | 4 | unclear |
| 120 | D125 | deaf | 22 | male | 75.33 | left | junior | 0 | severe | never | 8 | 2 | unclear |
| 121 | D126 | deaf | 23 | male | 76.68 | right | junior | 0 | severe | never | 5 | 2 | sudden |
| 122 | D127 | deaf | 19 | female | 79.26 | right | high | 0 | severe | never | 5 | 3 | sudden |
| 123 | D128 | deaf | 24 | female | 75.33 | right | junior | 0 | severe | never | 5 | 1 | sudden |
| 124 | D129 | deaf | 27 | female | 95.22 | right | college | 0 | severe | never | 6 | 3 | unclear |
| 125 | D130 | deaf | 24 | female | 75.33 | right | high | 0 | profound | never | 8 | 1 | sudden |

| | | | | | | | | | | | | | |
|---|---|---|---|---|---|---|---|---|---|---|---|---|---|
| 126 | D131 | deaf | 23 | female | 85.98 | right | college | 0 | severe | never | 7 | 1 | unclear |
| 127 | D132 | deaf | 24 | male | 102.36 | right | college | 0 | severe | never | 5 | 3 | sudden |
| 128 | D133 | deaf | 24 | male | 75.33 | right | primary | 2 | profound | never | / | / | unclear |
| 129 | D134 | deaf | 26 | male | 75.33 | right | junior | 4 | profound | never | 8 | 5 | heredity |
| 130 | D135 | deaf | 18 | male | 75.33 | left | college | 0 | severe | never | 3 | 3 | unclear |
| 131 | D136 | deaf | 18 | male | 82.48 | left | junior | 5 | profound | CI | 10 | 5 | unclear |
| 132 | D137 | deaf | 21 | male | 87.38 | left | high | 3 | severe | never | 10 | 2 | sudden |
| 133 | D138 | deaf | 24 | female | 75.33 | left | college | 3 | profound | everyday | 5 | 3 | sudden |
| 134 | D139 | deaf | 20 | male | / | right | college | 1 | profound | never | 7 | 2 | sudden |
| 135 | D140 | deaf | 34 | male | 78.7 | right | high | 0 | severe | never | 5 | 3 | sudden |
| 136 | D141 | deaf | 25 | male | 75.33 | right | college | 0 | profound | never | 7 | 3 | unclear |
| 1 | H001 | hearing | 23 | female | 124.67 | right | college | | | | | | |
| 2 | H002 | hearing | 21 | female | 124.67 | right | college | | | | | | |
| 3 | H003 | hearing | 24 | female | / | right | college | | | | | | |
| 4 | H004 | hearing | 20 | female | / | right | college | | | | | | |
| 5 | H005 | hearing | 21 | male | 114.02 | right | college | | | | | | |
| 6 | H006 | hearing | 30 | male | / | | college | | | | | | |
| 7 | H007 | hearing | 21 | female | / | right | college | | | | | | |
| 8 | H008 | hearing | 18 | male | 110.12 | right | college | | | | | | |
| 9 | H009 | hearing | 24 | male | / | right | college | | | | | | |
| 10 | H010 | hearing | 28 | female | / | right | college | | | | | | |
| 11 | H011 | hearing | 22 | female | 100 | right | college | | | | | | |
| 12 | H012 | hearing | 22 | female | 114.02 | right | college | | | | | | |
| 13 | H013 | hearing | 23 | female | 124.67 | right | college | | | | | | |
| 14 | H014 | hearing | 20 | male | 95.22 | right | college | | | | | | |
| 15 | H015 | hearing | 22 | female | 93.54 | right | college | | | | | | |
| 16 | H016 | hearing | 24 | male | 104.78 | right | college | | | | | | |
| 17 | H017 | hearing | 19 | female | / | right | college | | | | | | |
| 18 | H018 | hearing | 21 | female | 96.84 | right | college | | | | | | |
| 19 | H019 | hearing | 20 | male | 124.67 | right | college | | | | | | |
| 20 | H020 | hearing | 21 | male | / | right | college | | | | | | |
| 21 | H021 | hearing | 20 | female | 104.77 | right | college | | | | | | |

| | | | | | | | |
|---|---|---|---|---|---|---|---|
| 22 | H022 | hearing | 23 | female | 102.36 | right | college |
| 23 | H024 | hearing | 24 | female | 124.67 | right | college |
| 24 | H025 | hearing | 19 | female | / | right | college |
| 25 | H026 | hearing | 26 | male | 107.33 | right | college |
| 26 | H027 | hearing | 20 | female | 114.02 | right | college |
| 27 | H028 | hearing | 21 | female | 98.43 | right | college |
| 28 | H029 | hearing | 21 | male | 102.36 | right | college |
| 29 | H030 | hearing | 26 | female | 124.67 | right | college |
| 30 | H031 | hearing | 18 | female | 82.74 | right | college |
| 31 | H032 | hearing | 19 | female | 114.02 | right | college |
| 32 | H033 | hearing | 20 | female | 104.78 | right | college |
| 33 | H034 | hearing | 19 | female | 114.02 | right | college |
| 34 | H035 | hearing | 19 | male | 83.99 | right | college |
| 35 | H036 | hearing | 19 | female | 100 | right | college |
| 36 | H037 | hearing | 22 | female | 110.12 | right | college |
| 37 | H038 | hearing | 23 | female | 124.67 | right | college |
| 38 | H039 | hearing | 24 | female | 114.02 | right | college |
| 39 | H040 | hearing | 23 | female | / | right | college |
| 40 | H041 | hearing | 24 | female | 119.22 | right | college |
| 41 | H042 | hearing | 23 | female | 119.22 | right | college |
| 42 | H043 | hearing | 19 | female | 87.75 | right | college |
| 43 | H044 | hearing | 20 | male | 106.46 | right | college |
| 44 | H045 | hearing | 25 | male | 124.67 | left | college |
| 45 | H046 | hearing | 20 | male | 124.67 | right | college |
| 46 | H047 | hearing | 24 | male | 114.02 | right | college |
| 47 | H048 | hearing | 25 | male | 124.67 | right | college |
| 48 | H049 | hearing | 19 | female | 124.67 | right | college |
| 49 | H050 | hearing | 19 | female | 124.67 | right | college |
| 50 | H051 | hearing | 24 | female | 114.02 | right | college |
| 51 | H052 | hearing | 24 | female | 110.12 | right | college |
| 52 | H053 | hearing | 21 | female | / | right | college |
| 53 | H054 | hearing | 19 | female | 124.67 | right | college |

| | | | | | | | |
|---|---|---|---|---|---|---|---|
| 54 | H055 | hearing | 19 | female | / | right | college |
| 55 | H056 | hearing | 20 | male | 110.12 | right | college |
| 56 | H057 | hearing | 24 | female | 102.36 | right | college |
| 57 | H058 | hearing | 21 | female | 96.84 | right | college |
| 58 | H059 | hearing | 19 | male | 124.67 | right | college |
| 59 | H060 | hearing | 23 | female | 124.67 | right | college |
| 60 | H061 | hearing | 20 | male | 124.67 | right | college |
| 61 | H062 | hearing | 23 | female | 96.84 | right | college |
| 62 | H063 | hearing | 20 | male | 100 | right | college |
| 63 | H064 | hearing | 19 | male | 106.46 | right | college |
| 64 | H065 | hearing | 19 | female | 119.22 | right | college |
| 65 | H066 | hearing | 18 | female | / | right | college |
| 66 | H067 | hearing | 19 | male | 107.33 | right | college |
| 67 | H068 | hearing | 22 | male | 114.02 | right | college |
| 68 | H069 | hearing | 23 | male | / | right | college |
| 69 | H070 | hearing | 20 | male | 119.22 | right | college |
| 70 | H071 | hearing | 24 | male | 82.48 | right | college |
| 71 | H072 | hearing | 20 | male | 124.67 | right | college |
| 72 | H073 | hearing | 23 | male | 124.67 | right | college |
| 73 | H074 | hearing | 23 | male | 114.02 | right | college |
| 74 | H075 | hearing | 23 | male | 124.67 | right | college |
| 75 | H076 | hearing | 23 | male | 93.54 | right | college |
| 76 | H077 | hearing | 23 | male | 110.12 | right | college |
| 77 | H078 | hearing | 21 | female | 114.02 | right | college |
| 78 | H079 | hearing | 18 | male | / | right | college |
| 79 | H080 | hearing | 19 | female | 106.46 | right | college |
| 80 | H081 | hearing | 21 | female | / | right | college |
| 81 | H082 | hearing | 21 | female | 114.02 | right | college |
| 82 | H083 | hearing | 22 | female | 96.84 | right | college |
| 83 | H084 | hearing | 25 | female | / | right | college |
| 84 | H085 | hearing | 19 | female | 114.02 | right | college |
| 85 | H086 | hearing | 21 | female | 110.12 | right | college |

| | | | | | | | |
|---|---|---|---|---|---|---|---|
| 86 | H087 | hearing | 20 | female | 124.67 | left | college |
| 87 | H088 | hearing | 20 | female | / | right | college |
| 88 | H089 | hearing | 22 | female | / | right | college |
| 89 | H090 | hearing | 19 | female | / | right | college |
| 90 | H091 | hearing | 20 | male | 77.48 | right | college |
| 91 | H092 | hearing | 19 | male | 95.22 | right | college |
| 92 | H093 | hearing | 19 | female | 119.22 | right | college |
| 93 | H094 | hearing | 19 | female | 119.22 | right | college |
| 94 | H095 | hearing | 19 | female | 95.22 | right | college |
| 95 | H096 | hearing | 19 | female | 124.67 | right | college |
| 96 | H097 | hearing | 23 | male | 114.02 | right | college |
| 97 | H098 | hearing | 19 | male | 103.16 | right | college |
| 98 | H099 | hearing | 19 | male | 80.78 | right | college |
| 99 | H100 | hearing | 22 | male | 114.02 | right | college |
| 100 | H101 | hearing | 22 | female | 124.67 | right | college |
| 101 | H102 | hearing | 19 | female | 106.46 | right | college |
| 102 | H103 | hearing | 21 | female | 124.67 | right | college |
| 103 | H104 | hearing | 23 | female | 102.36 | right | college |
| 104 | H105 | hearing | 25 | male | 114.02 | right | college |
| 105 | H106 | hearing | 19 | female | 100 | right | college |
| 106 | H107 | hearing | 19 | female | 124.67 | right | college |
| 107 | H108 | hearing | 18 | female | 124.67 | right | college |
| 108 | H109 | hearing | 24 | female | 114.02 | right | college |
| 109 | H110 | hearing | 26 | female | 104.78 | right | college |
| 110 | H111 | hearing | 24 | male | 124.67 | right | college |
| 111 | H112 | hearing | 22 | female | / | right | college |
| 112 | H113 | hearing | 21 | female | 107.33 | right | college |
| 113 | H114 | hearing | 25 | male | 104.78 | right | college |
| 114 | H115 | hearing | 22 | female | / | right | college |
| 115 | H116 | hearing | 21 | female | 119.22 | right | college |
| 116 | H117 | hearing | 21 | male | 98.43 | right | college |
| 117 | H118 | hearing | 21 | female | 98.43 | right | college |

| | | | | | | | |
|---|---|---|---|---|---|---|---|
| 118 | H119 | hearing | 22 | male | 107.33 | right | college |
| 119 | H120 | hearing | 21 | female | 104.78 | right | college |
| 120 | H121 | hearing | 19 | female | 110.12 | right | college |
| 121 | H122 | hearing | 22 | female | 114.02 | right | college |
| 122 | H123 | hearing | 23 | male | / | right | college |
| 123 | H124 | hearing | 24 | female | 104.78 | right | college |
| 124 | H125 | hearing | 23 | female | 98.43 | right | college |
| 125 | H126 | hearing | 22 | female | 85.98 | right | college |
| 126 | H127 | hearing | 22 | male | 75.33 | right | college |
| 127 | H128 | hearing | 21 | male | 114.02 | right | college |
| 128 | H129 | hearing | 21 | male | 102.36 | right | college |
| 129 | H130 | hearing | 21 | male | 96.84 | right | college |
| 130 | H131 | hearing | 24 | male | / | right | college |
| 131 | H132 | hearing | 23 | male | 96.84 | right | college |
| 132 | H133 | hearing | 20 | male | 107.33 | right | college |
| 133 | H134 | hearing | 22 | male | 119.22 | right | college |
| 134 | H135 | hearing | 25 | male | / | right | college |
| 135 | H136 | hearing | 20 | female | 95.22 | right | college |

Note: Participants shown with gray shading were excluded from formal data analyses; CI = cochlear implantation; "/" indicates that the information was not provided

**Table S7. Sample size in each task**

| task | session | group | number of total participants | number of eligible participants |
|---|---|---|---|---|
| **identity** | dynamic | hearing | 134 | 131 |
| | | deaf | 132 | 128 |
| | static | hearing | 113 | 113 |
| | | deaf | 104 | 100 |
| **emotion** | dynamic | hearing | 135 | 135 |
| | | deaf | 134 | 129 |
| | static | hearing | 134 | 134 |
| | | deaf | 133 | 129 |
| **speech** | normal | hearing | 134 | 134 |
| | | deaf | 133 | 129 |
| | reverse | hearing | 116 | 116 |
| | | deaf | 106 | 103 |
| **global motion** | dynamic | hearing | 128 | 128 |
| | | deaf | 124 | 122 |

Note: Because not all participants completed every session, the sample size varied across sessions. For each linear mixed-effects analysis, we included only the participants who completed all sessions of the task.
